# Multidimensional Scaling and Relatedness Research

**DOI:** 10.1101/297879

**Authors:** Jan Graffelman, Iván Galván Femenía, Rafael de Cid, Carles Barceló-i-Vidal

**Affiliations:** Department of Statistics and Operations Research Universitat Politècnica de Catalunya, Barcelona, Spain; Department of Biostatistics University of Washington, Seattle, USA; Department of Computer Science, Applied Mathematics & Statistics Universitat de Girona, Girona, Spain; Genomes For Life - GCAT lab Program of Predictive and Personalized Medicine of Cancer (PMPPC) Institute for Health Science Research Germans Trias i Pujol (IGTP) Can Ruti Campus, Badalona, Barcelona, Spain

**Keywords:** allele sharing, composition, genetic bootstrapping, identity by state, identity by descent, log-ratio transformation

## Abstract

Multidimensional scaling is a well-known multivariate technique, that is often used in genetics for studying population substructure. In this paper we show that multidimensional scaling of marker data is of relevance for relatedness research. Relatedness is usually investigated by estimating and plotting identity-by-state and identity-by-descent allele-sharing statistics. We show that outlying individuals in a map obtained by multidimensional scaling of genetic variables do not necessarily stem from a different human population, but can be the consequence of relatedness. We propose a method for classifying pairs of individuals into the standard relationship categories that combines genetic bootstrapping, multidimensional scaling and discriminant analysis. We validate our method with simulation studies. Given the variant filtering procedures, our method classifies relationships up to and including the fourth degree with high accuracy (96-97%), using only identity by state. The usefulness of the method is illustrated with data from the 1,000 genomes and the GCAT projects.

## 1 Introduction

Multidimensional scaling (MDS) is a versatile multivariate statistical technique that has found applications in many fields of science [Cox and Cox, 2001, Borg and Groenen, 2005]. In genetics, MDS is often used to investigate the existence of population substructure. If individuals do not come from a single homogeneous human population, but from two or more populations with different allele frequencies, then an MDS of the genetic marker data typically separates the individuals of the different populations. An example is given in Figure 1 where we show the MDS map for a sample of 368 individuals, 165 from the CEU population, and 203 from the YRI population of the HapMap project [The International HapMap Consortium, 2007], now included in the 1,000 genomes project [The 1000 Genomes Project Consortium, 2015], using the allele-sharing distance [Mountain and Cavalli-Sforza, 1997]. Many examples of the use of MDS for detecting population substructure can be found in the genetics literature. Pemberton *et al.* (2010) used MDS to investigate the population affiliation of the HapMap Phase III individuals. Other examples of population substructure research with MDS are given by Jakobsson *et al.* (2008), Sabatti (2009), Foulkes (2009), Wang (2010) and Pemberton *et al.* (2013).

**Figure 1:**
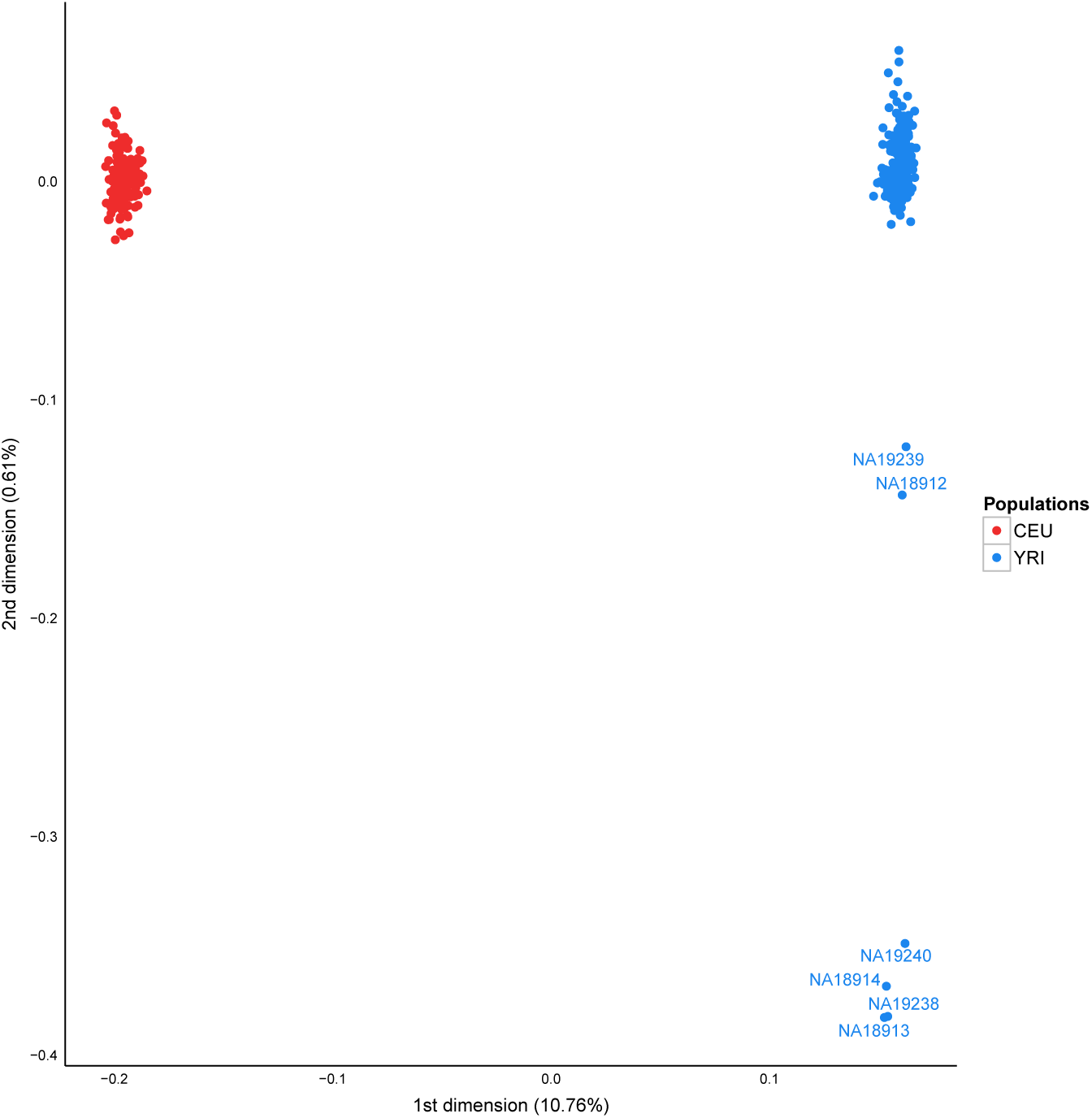
MDS map for a sample of 165 individual of the CEU and 203 individuals of the YRI population of the 1,000 genomes project, using 48,072 LD-pruned, autosomal SNPs with MAF *>* 0.40 and non-significant in a HWE exact test.

The examples in the literature and in Figure 1 suggest that observing outlying individuals in an MDS map implies these individuals are different, and therefore potentially stem from a different human populations that may be characterized by different allele frequencies for the variants under study. In this paper we show that this conclusion is not always warranted. Outlying individuals, and even groups of outlying individuals, may very well stem from the *same* human population and appear separated from their population for a very different reason: they correspond to individuals that are closely related. Figure 1 illustrates this point, where some interrelated individuals of the YRI population appear separated from their alikes. In particular, the map distance of the most outlying individuals with respect to the center of the YRI population is actually *larger* than the distance between the centers of the YRI and the CEU population. The four most outlying individuals are all involved in two parent-offspring (PO) relationships, whereas the two intermediate outliers are involved in one PO relationship (Pemberton *et al.* (2010), supplemental data). Relatedness investigations in genetics are mostly based on allele-sharing statistics such as the average number of IBS alleles shared by a pair of individuals over a set of loci, or by estimating the probabilities of sharing 0, 1 or 2 alleles IBD [Cotterman, 1941, Thompson, 1991], known as Cotterman’s coefficients. Plots of these sharing statistics typically show clusters that correspond to unrelated individuals (UN), parent-offspring pairs (PO), full sibs (FS), half sibs (HS), monozygotic twins (MZ), avuncular pairs (AV), first cousins (FC), grandparent-grandchild (GG) or more remote relationships. MDS is not used in such relatedness investigation, but is linked to the allele sharing approach in the sense that a distance matrix can easily be computed from the allele sharing statistics. Moreover, recent studies [Weir and Goudet, 2017, Conomos et al., 2016] have pointed out that relatedness and population substructure are tightly connected topics in the sense that both induce allele sharing.

The remainder of this paper is organized as follows. In Section 2 we provide background on relatedness research and MDS, and we develop theory for using MDS in relatedness research by calculating genetic distances between pairs of individuals. Section 3 explores and validates our methodology with some simulation studies, that assess the effects of sample size, number of variants and minor allele frequency (MAF) on the classification error rate obtained by a discriminant analysis based on MDS results. Section 4 shows applications of the proposed methodology with data from the 1,000 genomes and GCAT projects. A Discussion section (5) finishes the paper.

## 2 Theory

### Relatedness research

We briefly review some fairly standard procedures that are currently used in relatedness research. Relatedness investigations are focused on the extent to which alleles are shared between individuals. Two individuals can share 0, 1 or 2 alleles for any autosomal variant. Alleles can be identical by state (IBS) or identical by descent (IBD). A pair of individuals share IBS alleles if they match irrespective of their provenance; whereas they share IBD alleles only if they come from a common ancestor. In IBS studies, the means and standard deviations of the IBS allele counts over variants [Abecasis et al., 2001], or the proportions of variants sharing 0, 1 and 2 IBS alleles [Rosenberg, 2006] can be plotted. These plots reveal characteristic clusters corresponding to MZ, PO, FS, HS/AV/GG, FC or UN pairs. Alternatively, in an IBD based approach, the probability of sharing 0, 1 or 2 IBD alleles (usually denoted by 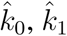 and 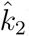) can be represented in a scatterplot [Nembot-Simo and Graham, 2013]. The Cotterman coefficients can be estimated by the method of moments [Purcell et al., 2007] or by maximum likelihood [Thompson, 1975, Milligan, 2003, Weir et al., 2006]. A good account of the state-of-the-art of IBD relatedness research is given by Weir *et al.* (2006). In IBD studies, reference values for the standard relationships are available (see Table 1). Related pairs can also be distinguished, albeit at lower resolution, by using the coancestry coefficient defined as *θ* = *k*_1_*/*2 + *k*_2_ or the kinship coefficient defined as *φ* = *θ/*2. An example of the use of *φ* is shown by Conomos *et al.* (2016) where plots of the kinship coefficient and *k*_0_ also uncover clusters with the different family relationships. Galvaán *et al.* (2017) give an overview of graphics for related ness research. Figure 2 shows a panel plot of some standard graphics used in IBS and IBD studies for all the pairs of individuals from the CEU population of the HapMap project. These plots distinguish UN, PO, FS and second degree pairs. First and second degree relationships for the CEU population were documented by Pemberton *et al.* (2010) using IBS methods, and confirmed by Kyriazopoulou *et al.* (2011), who used hidden Markov models and suggested additional third and fourth degree relationships. Stevens *et al.* (2012) used IBD methods confirming the results of Pemberton *et al.* (2010).

**Table 1:** Degree of relationship (R), kinship coefficient (*φ*), and probability of sharing zero, one or two alleles identical by descent (*k*_0_, *k*_1_, *k*_2_).

| Type of Relative | R | $\phi$ | IBD Probabilities | | |
| --- | --- | --- | --- | --- | --- |
| | | | $k_0$ | $k_1$ | $k_2$ |
| Monozygotic twins (MZ) | 0 | 1/2 | 0 | 0 | 1 |
| Full-siblings (FS) | 1 | 1/4 | 1/4 | 1/2 | 1/4 |
| Parent-offspring (PO) | 1 | 1/4 | 0 | 1 | 0 |
| Half-siblings/ grandchild-grandparent/<br>niece/nephew-uncle/aunt (HS,GG,AV) | 2 | 1/8 | 1/2 | 1/2 | 0 |
| First cousins (FC) | 3 | 1/16 | 3/4 | 1/4 | 0 |
| Unrelated (UN) | ∞ | 0 | 1 | 0 | 0 |

**Figure 2:**
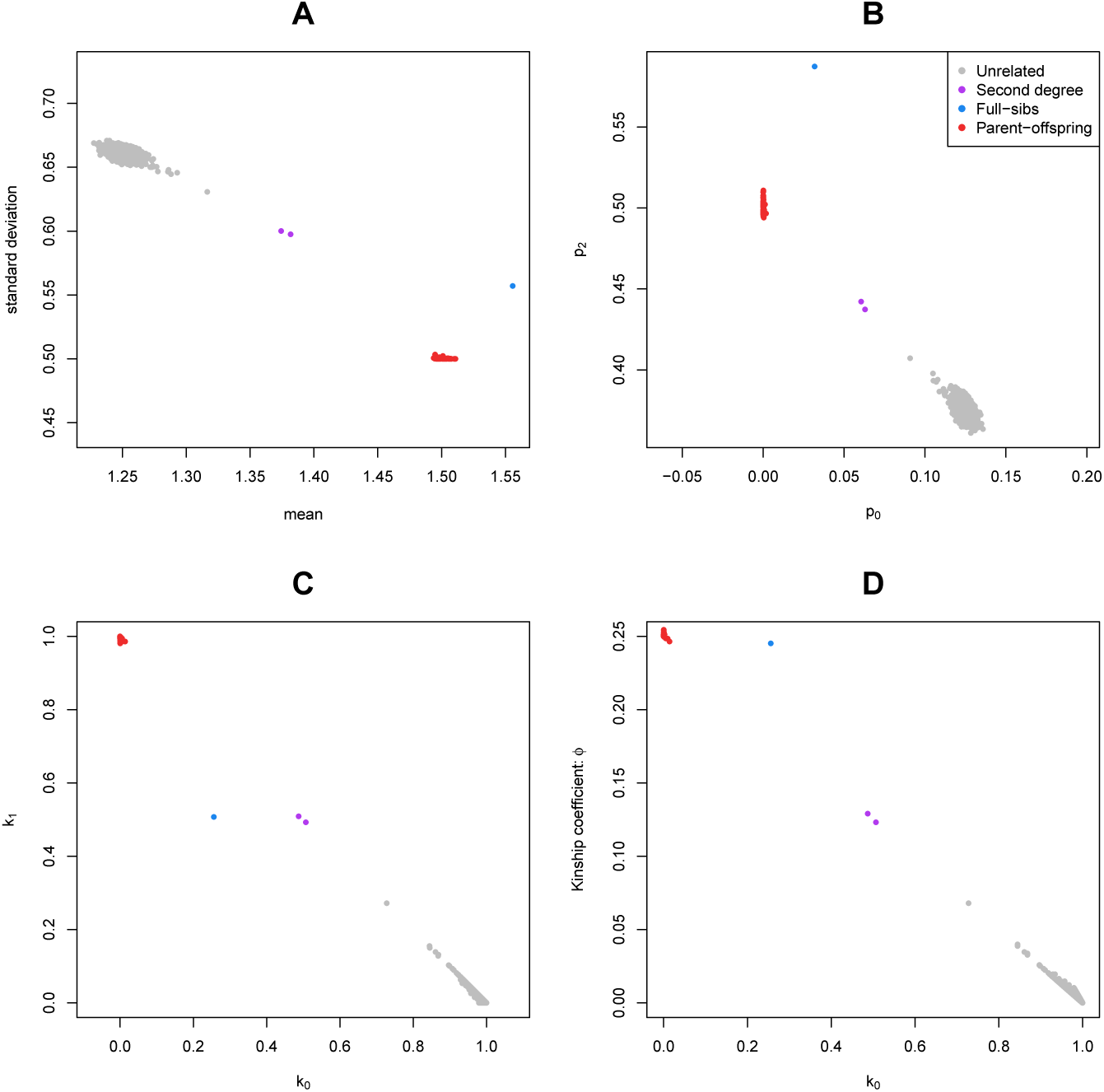
Some standard graphics for relatedness research for the CEU sample of the HapMap project. IBS/IBD statistics were calculated over a set of 26,081 complete, LD-pruned autosomal SNPs with MAF above 0.4, and HWE exact test p-value above 0.05. A: scatterplot of the mean and standard deviation of the number of IBS alleles. B: scatterplot of the fraction of variants sharing 2 (*p*_2_) against the fraction sharing 0 (*p*_0_) IBS alleles. C: scatterplot of 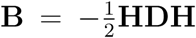 against 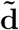. D: scatterplot of *φ* against 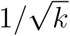.

Alternatively, a Markov-chain approach with the calculation of likelihood ratios for putative and alternative relationship has been developed by Epstein *et al.* (2000) (the Relpair program) and by McPeek and Sun (2000) (the Prest-plus program).

### Multidimensional scaling and relatedness research

Multidimensional scaling (MDS) is a well-known multivariate method with many applications in genetics, though its potential in relatedness research seems not to have been explored. Several variants of multidimensional scaling exist and a distinction is usually drawn between metric and non-metric multidimensional scaling. Metric MDS is also known as classical scaling or principal coordinate analysis (PCO, Gower [1966]). In metric MDS a map of the individuals is obtained by extracting eigenvalues and eigenvectors from a conveniently transformed distance matrix. The distances in the map approximate the (genetic) distances between the individuals. The method is not invariant under monotone transformations of the original distances. Non-metric multidimensional scaling obtains the map in an iterative manner by minimizing a stress function, and is invariant under monotone transformations of the distances, as it uses only the rank order of the distances. In this paper we confine ourselves mostly to metric MDS. The theory of MDS is described in most textbooks on multivariate analysis [Mardia et al., 1979] and some books are entirely dedicated to the topic [Cox and Cox, 2001, Borg and Groenen, 2005].

Figure 3 shows metric MDS maps for simulated populations with some related individuals. Bi-allelic marker data was generated by sampling independent genotypes from a multinomial distribution with Hardy-Weinberg frequencies, using uniformly distributed allele frequencies. Some related individuals were then generated from the base set of independent individuals. Figure 3A shows that a single monozygotic twin in the database stands out as an outlier in the map (the two twin-sibs show up as two coinciding points in the first dimension). The existence of two identical points in MDS can also be inferred from the eigenvalues, as two identical points create an additional zero eigenvalue in the MDS output (one zero eigenvalue is always present and is a characteristic of the method as it results from the double-centring of the transformed distance matrix). At first sight the appearance of related individuals as outliers seems contradictory. The MZ pair in panel (A) is shown at a large distance from the rest of the population suggesting that the genetic distance between a twin-sib and the rest of the individuals is much larger than any of the distances between any other pair of unrelated non-twin-sib individuals. This is actually not true. The genetic distances between a twin-sib and the other individuals are in fact of similar order of magnitude as the distances between any pair of unrelated individuals. The reason that the MZ pair separates out in the map at a larger distance is due to the fact that the MZ pair constitutes an outlier in the analysis. The genetic distance between the twin-sibs is zero, and this is an extreme outlier in the off-diagonal entries of the distance matrix, where all distances between unrelated pairs are much larger. MDS is sensitive to the existence of outliers and fits the outlying point much better than the other points. In regression, it is well-known that a single outlying observation can exert leverage on the slope of the regression line and can highly influence the slope [Weisberg, 2005, Chapter 9]. This phenomenon also arises in MDS, where a single outlying observation exerts leverage on the fitted plane. Consequently, the goodness-of-fit of all distances involving the twins are all high, whereas the goodness-of-fit of all other distances is very low, as is revealed in a diagnostic plot of fitted versus observed distances (see supplementary Figure S1). Similarly, most of the maps shown in Figure 3 containing some related individuals are pretty much uninformative on most of the distances, except for those distances that involve the outlying points.

**Figure 3:**
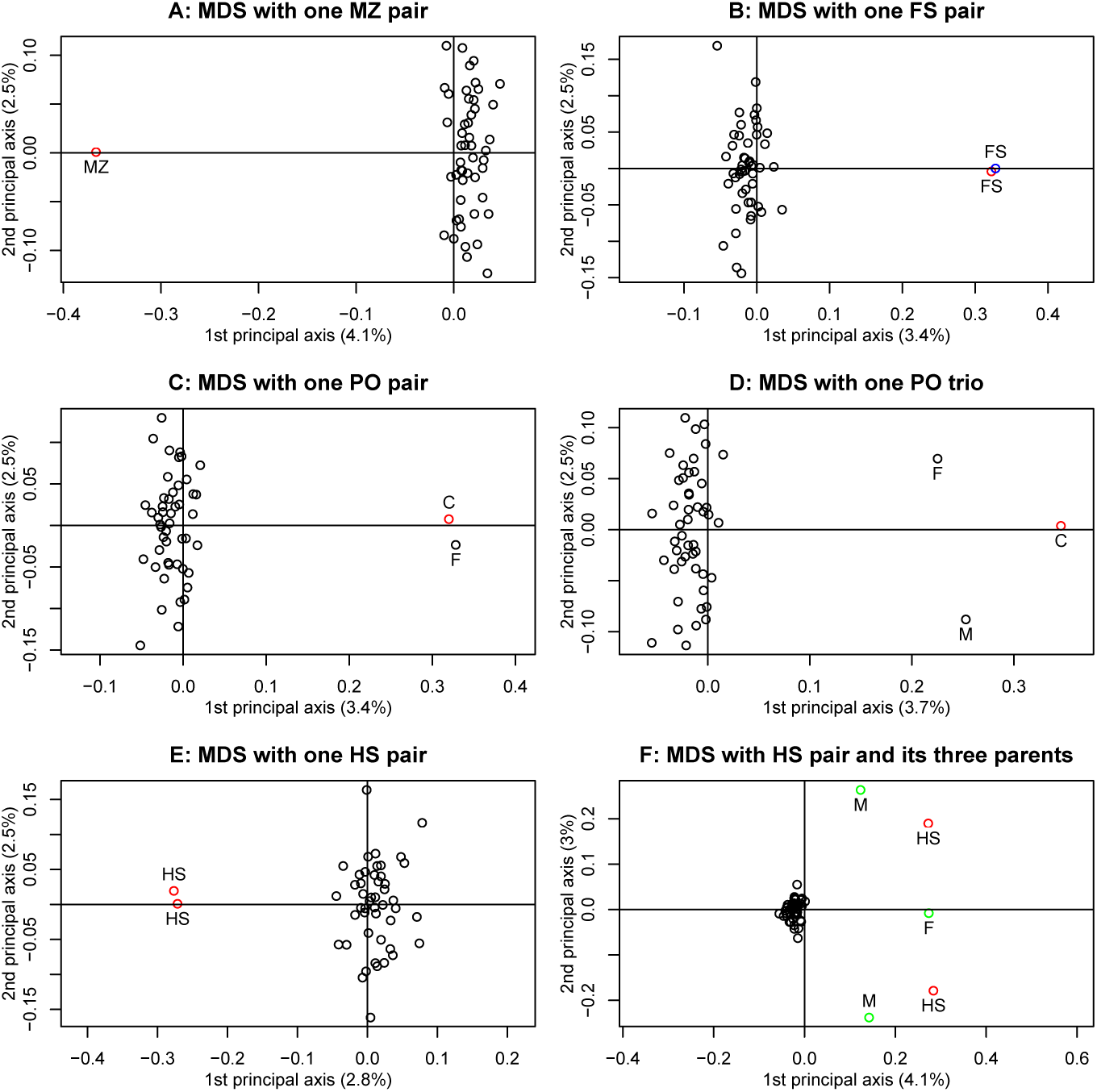
MDS maps for simulated populations with some related individuals. (F=Father, M=Mother, C=Child, MZ=Monozygotic twin, FS=Full Sib, HS=Half Sibs)

**Figure 4:**
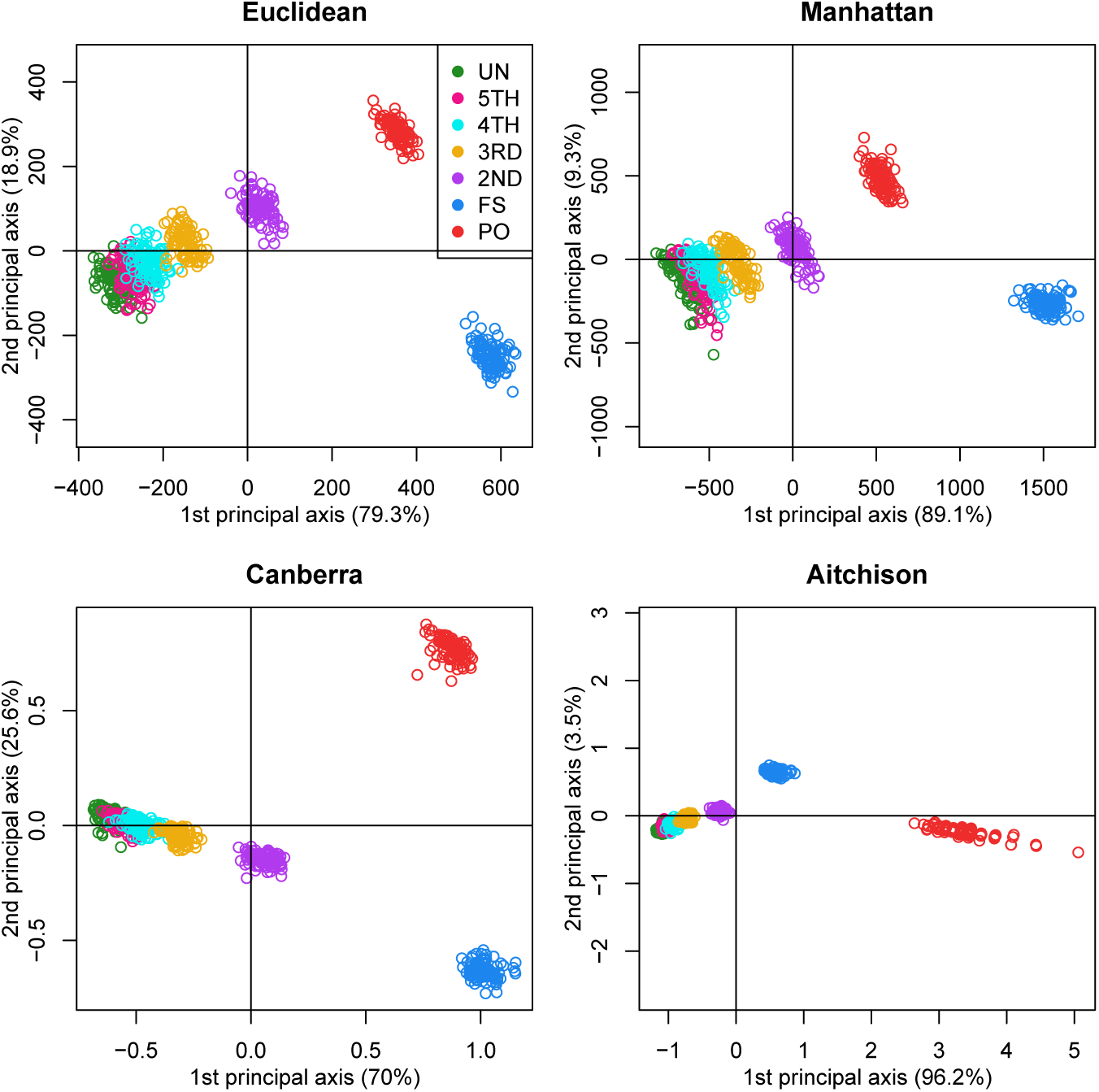
MDS maps for simulated samples, with 1% genotyping error. 100 pairs of each type of relationship (UN, fifth, fourth, third (FC), second (HS), FS and PO) were generated using an allele frequencies of 0.5, assuming Hardy-Weinberg equilibrium and 5,000 independent bi-allelic variants.

### MDS with pairs of individuals

An important difference between the standard allele-sharing methods outlined in the first paragraph of this Section and MDS as discussed above is that the allele-sharing methods plot *pairs* of individuals, whereas MDS is usually employed to plot the *individuals themselves*. To bring MDS maps to the same realm as the allele sharing techniques, we propose to compute a distance matrix between all possible *pairs* of individuals rather than the distance matrix between the individuals. In order to do so, we need to define a distance (or similarity) measure between two pairs of individuals. This can be done in many different ways. The distance measure needs to indicate to what extend two pairs of individuals are similar or not. We use *n* for the number of individuals, and *k* for the number of genetic variants. For bi-allelic variants there exist six possible pairs of genotypes whose counts over *k* variants can be laid out in a triangular array shown in Table 2, where *k_ij_* refers to the number of variants that have *i* B alleles for one individual, and *j* B alleles for the other individual.

**Table 2:** Lower triangular matrix layout with counts for all possible genotype pairs.

|  |  |  |  |  |
| --- | --- | --- | --- | --- |
| 1st indiv. | AA | $k_{00}$ | | |
| | AB | $k_{10}$ | $k_{11}$ | |
| | BB | $k_{20}$ | $k_{21}$ | $k_{22}$ |
|  |  | AA | AB | BB |
| 2nd indiv. |  |  |  |  |

Consequently, each pair can be represented by a vector of six counts which can be expressed as a composition by division by its total 

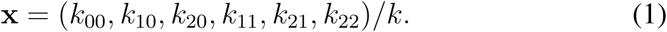

The total number of variants is given by *k* = Σ_*i*≥*j*_ k_*ij*_. For PO pairs this vector has, in theory, a structural zero, *k*_20_ = 0, because PO pairs share at least one IBS allele. However, for empirical data *k*_20_ = 0 is, with large *k*, never observed due to the existence of some mutations and genotyping error. In order to measure the distance between two pairs, **x** and **y**, we consider the ordinary Euclidean distance (*d_e_*), the Manhattan distance (*d_m_*), the Canberra distance (*d_c_*) and the Aitchison distance [Aitchison, 1986], *d_a_*, where the latter is appealing for being specifically designed for compositional data. These distances are given by 

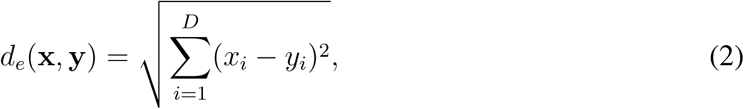

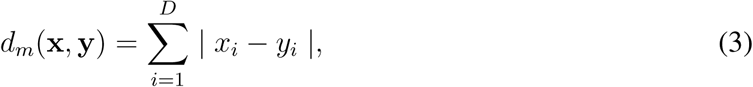

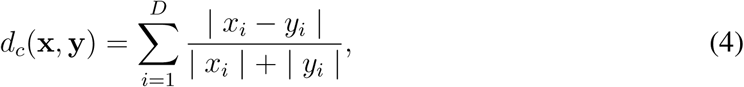

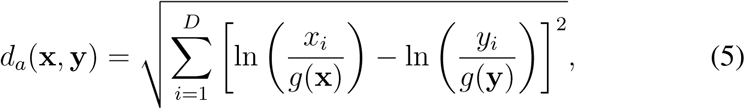

where *D* represents the number of parts of the composition (here *D* = 6), and *g*(*•*) the geometric mean of the composition. The Aitchison distance requires strictly positive elements, which is theoretically a problem for PO pairs. Methodology for dealing with zeros in compositions has been developed (Mart´ın-Fernaändez *et al.*, 2011). Additional metrics could be explored, but are not considered here.

A drawback of the representation of pairs of individuals in an MDS map is that the type of relationship cannot be inferred. Outlying points observed in pairwise MDS maps are likely to correspond to related pairs of individuals, but without additional analysis one does not know whether the outlier is an MZ, PO, FS, HS or other pair. We resolve this by first identifying a subset of approximately unrelated individuals in the database, having a co-ancestry coefficient with other individuals that is below 0.05. We next simulate pairs of related individuals of known relationships from this subset, applying the Mendelian inheritance rules. E.g., PO pairs are simulated by first drawing two parents at random from the unrelated subset. A child is then simulated by drawing one allele at random from both these parents. The process is repeated in order to generate many random PO pairs. FS, HS and pairs of other relationships are simulated in an analogous manner. This process is based on re-sampling the alleles of the individuals, and we will refer to this principle as *genetic bootstrapping*. The artificially generated data set forms a *reference set* or *training set* against which the empirically observed data can be compared. This reference set is conditional on the allele frequencies of the observed sample. The distance matrix of all simulated pairs is calculated and used to construct an MDS reference map. The empirically observed pairs are projected onto this MDS map and their relationship is inferred, according to which simulated type of relationship is most close to the empirical pair. This can be done in a quantitative way by classifying all empirical pairs with linear discriminant analysis, using the simulated pairs as a training set. We briefly outline the calculations below.

MDS [Mardia et al., 1979, Chapter 14] is performed by the spectral de composition in Equation (6) of the *m × m* scalar product matrix **B**, which is obtained from the double centred matrix with the squared distances (**D**) as 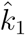, where **H** is an idempotent centring matrix (**H** = **I** − (1/*m*)**11**^′^), 
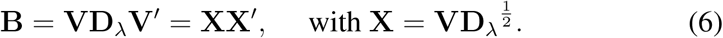

Matrix **V** contains the eigenvectors of **B**, and **D**_*λ*_ is a diagonal matrix containing the eigenvalues in non-increasing order of magnitude. Matrix **X** contains the coordinates of the pairs in an MDS map of dimension *r*, the dimensionality chosen for the representation of the results.

Let **d** be the *m ×* 1 vector of supplementary distances, containing the squared distances between an empirical pair and all *m* pairs in the reference set. The squared transformed distances 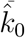 are obtained as 

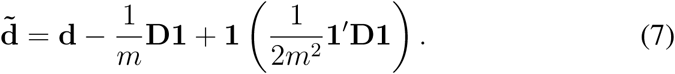

This transformation amounts to subtracting the average of the squared distance vector and adding half the mean of the squared entries of the original distance matrix calculated for the reference set. The projection of the empirical pairs is carried out by using results that plot supplementary points in an MDS map. In particular, the coordinates of a projected pair, **z**, are found by [Gower, 1968] 

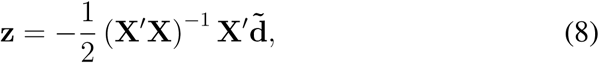

where the resulting coordinate vector **z** is *r ×* 1, and has as many elements as the dimensionality chosen for representation of the results. The validity of (7) and (8) is easily verified by taking a copy of one row of the original distance matrix, and applying the projection equations. The map coordinates of the projected point will then coincide exactly with the coordinates of that point in the initial MDS solution. Equations (7) and (8) exemplify that a supplementary point is found by first centring (and scaling) the observation in the same way the original observations are centered (and scaled), followed by a regression [Graffelman and Aluja-Banet, 2003].

## 3 Simulations

We first validate the proposed methodology with some simulations. We simulated 5,000 independent genetic bi-allelic variants by sampling from a multinomial distribution under the Hardy-Weinberg assumption, using a minor allele frequency of 0.5 for all variants. Using Mendelian inheritance rules, 100 independent pairs of each type of relationship were simulated. We used two scenarios, one with perfect application of Mendelian inheritance rules, yielding simulated data sets that are free of Mendelian inconsistencies, and one with 1% genotyping error for all three genotypes. Two-dimensional metric MDS maps using four distance measures (Euclidean, Manhattan, Canberra and Aitchison) for the six-part compositions of the pairs are shown in Figure 4. An MDS with the Euclidean distance is seen to separate the main types of relationships until the fourth degree (UN, fourth, third, and second degree, FS and PO), with overlapping clusters for UN, fifth and fourth degree. The FS group is most outlying. The Manhattan metric gives a picture that is similar to the Euclidean distance, but with apparently less discrimination between UN, fifth, fourth, third and second degree relationships. The Canberra metric, which is related to the Manhattan metric, places PO and FS pairs at similar positions in the first dimension, and separates them in the second. Results obtained with the Aitchison metric differ, as this metric strongly sorts out the PO pairs in the first dimension, inverting the order of the PO and FS clusters in this dimension. Scatterplots of the MDS coordinates beyond the second dimension show, in general, little separation of the different relationships, confirming that the solution is essentially two-dimensional. Exceptions are the Canberra and Aitchison distance, which do show clear clustering of relationships in the third dimension (see supplementary figure S2). Classification rates for a linear discriminant analysis, using the first three MDS dimensions, are given for the four metrics in Table 3.

**Table 3:** Overall and up to and including fourth degree relationships classification rates for linear discriminant analysis using four different metrics, and applying genetic bootstrapping without genotyping error, or with 1% genotyping error.

|  |  | Metric |  |  |  |
| --- | --- | --- | --- | --- | --- |
|  | Genotyping error | Euclidean | Manhattan | Canberra | Aitchison |
| Overall | 0% | 0.910 | 0.899 | 0.914 | 0.915 |
|  | 1% | 0.910 | 0.884 | 0.906 | 0.851 |
| Up to 4th | 0% | 0.966 | 0.952 | 0.960 | 0.964 |
|  | 1% | 0.952 | 0.942 | 0.946 | 0.934 |

The non-perfect overall classification can mainly be ascribed to the fifth degree pairs, who are partly misclassified as fourth degree or unrelated. The classification rates up to and including the fourth degree pairs are about five percent higher. The differences between the metrics are small, though the Euclidean distance seems to work best, in particular in the presence of genotyping error. Somewhat better classification rates are obtained if genotyping error is absent. When there is no genotyping error, there are structural zeros in the compositions of PO pairs, which were dealt with using methods outlined by Martín-Fernández *et al.* (2011). Similar results were obtained by using quadratic discriminant analysis (not shown).

This simulation concerns a relatively ideal dataset with independent variants and maximally polymorphic variants. For empirical data sets, the independence of the variants can be approximately achieved by LD pruning variants. In practice, many variants have a low MAF. We investigated the effect of the MAF on the discriminatory power of the different metrics, by simulating variants with different MAF. Figure 5 shows how the classification rate varies as a function of the MAF and the number of bootstrap samples for the considered metrics.

**Figure 5:**
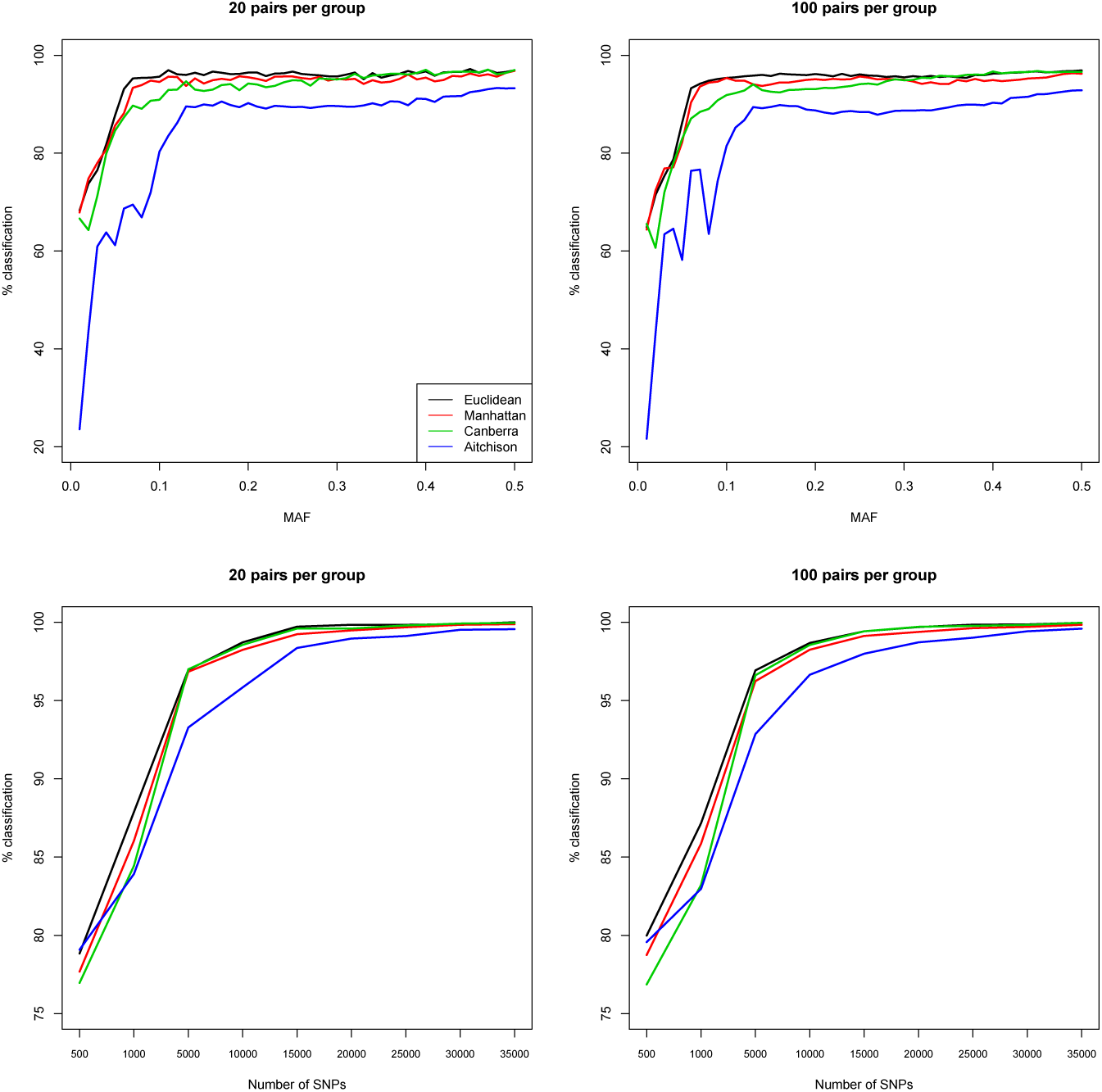
Classification rate up to and including the fourth degree as a function of the minor allele frequency (MAF) and the number of maximally polymorphic variants obtained by linear discriminant analysis using four different metrics, and allowing for 1% genotyping error.

Figure 5 shows a worse classification rate for the Aitchison metric, whereas Euclidean, Manhattan and Canberra metrics achieve the best classification with high MAF variants. The observed classification rates are similar for a training set with 20 and 100 simulated pairs. The classification rate depends on the number of maximally polymorphic variants, and suggests that 25,000 variants are sufficient to almost perfectly classify PO, FS, second, third and fourth degree relationships. In the light of the simulation results, we will use one hundred bootstrap samples and the Euclidean distance for the examples presented in Section 4. In the remainder of the paper, we will refer to our method, which combines genetic bootstrapping, IBS allele sharing, MDS and LDA, as IBS-MDS.

## 4 Applications

In this Section we apply the proposed MDS of pairwise distances to two genomic data sets. We use the CEU population of the HapMap project, now available as part of the 1,000 genomes project [The 1000 Genomes Project Consortium, 2015] (www.internationalgenome.org), whose family relationships have been analysed in detail by Pemberton *et al.* (2010), Kyriazopoulou *et al.* (2011), Huff *et al.* (2011) and Stevens *et al.* (2011, 2012). We also present a relatedness study of the population based GCAT Genomes for Life project (a cohort study of the genomes of Catalonia, www.genomesforlife.com, Obo´n-Santacana et al. [2018]). For both projects, we used Plink 1.90 [Purcell et al., 2007] for data manipulation and filtering, and R [R Core Team, 2014] for MDS and discriminant analysis.

### 4.1 The CEU sample

We detail the analysis of the CEU panel. HapMap phase II and III variants of the CEU panel were filtered according to missingness (only variants genotyped for all individuals were used), MAF (*>* 0.40) and Hardy-Weinberg equilibrium test result (using the exact mid p-value *>* 0.05 [Graffelman and Moreno, 2013]). Variants were LD-pruned with Plink’s independent pairwise option (indep-pairwise 50 5 0.2). The final data set contained 31,370 autosomal variants. The CEU panel consists of 165 individuals, mainly PO trios, giving 13,530 possible pairs of individuals. The ordinary MDS map of the individuals of this panel is shown in Figure 6. Documented related individuals are connected by lines. Figure 6 shows an outlying FS pair in the first dimension, that pulls the related families along this axis.

**Figure 6:**
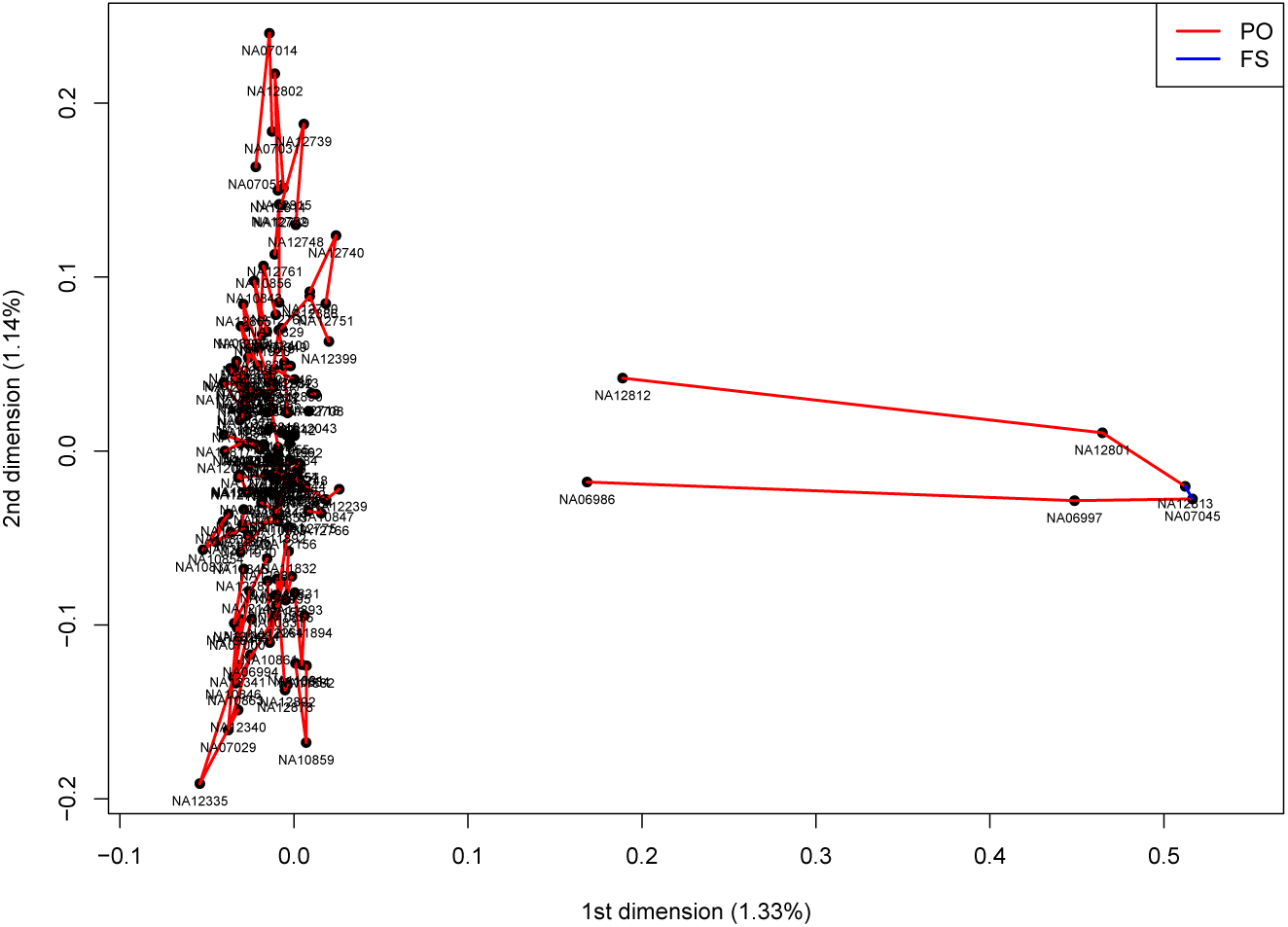
MDS map for a sample of 165 individuals of the CEU population, with outlying FS pair in the first dimension

Figure 7 shows the MDS map of the pairwise approach, using the Euclidean distance measure. The convex hulls delimit the cloud of the positions of the simulated UN, fifth, fourth, third, second, FS and PO pairs (using 100 pairs of each). The classification rate of the simulated data was as high as 99.6%. Points with open circles represent the empirical pairs which were projected onto the map using Equations (7) and (8), and are coloured according to their predicted relationship. The map suggests the CEU sample has 96 PO pairs, one FS pair, two second degree pairs, one third degree pair, five fourth degree pairs and many fifth degree pairs that merge with UN pairs. The classification of the empirical pairs by linear discriminant analysis confirmed the 96 PO and the single FS pair relationships described by Pemberton *et al.* (2010) (results not shown), as well as the additional FC pair reported by Kyriazopoulou *et al.* (2011), with a posterior probability of one for all these pairs. Third and fourth degree relationships uncovered by Kyriazopoulou *et al.* (2011) are reported in Table 4, together with the posterior probabilities obtained in our IBS-MDS analysis. 193 pairs were classified as fifth degree relationship pairs, of which 66 had a posterior probability above 0.95. We do not report all these pairs because given the overlap with the UN cluster and the poorer classification rate of the fifth degree observed in the simulations.

**Table 4:** Predicted relationships of third (3rd) and fourth (4th) degree pairs of the CEU sample of the HapMap project reported by Kyriazopoulou *et al.* (2011) and their posterior probabilities according to IBS-MDS. Coding and abbreviations used: sex 1 = male, 2 = female; a hyphen (-) indicates the corresponding pair is not annotated or regarded unknown by the corresponding authors; FCOR: first cousin once removed; Pem.: Pemberton *et al.* (2010); Kyr.: Kyriazopoulou *et al.* (2011); Ste.: Stevens *et al.* (2012); Huf.: Huff *et al.* (2011).

| Pair | ID1 | sex | ID2 | sex | Pem. | Kyr. | Ste. | Huf. | IBS-MDS | Posterior probabilities |  |  |  |  |  |  |
| --- | --- | --- | --- | --- | --- | --- | --- | --- | --- | --- | --- | --- | --- | --- | --- | --- |
|  |  |  |  |  |  |  |  |  |  | UN | 5th | 4th | 3rd | 2nd | FS | PO |
| 1 | NA06997 | 2 | NA12801 | 1 | - | FC | FC | - | 3rd | 0.000 | 0.000 | 0.000 | 1.000 | 0.000 | 0.000 | 0.000 |
| 2 | NA06993 | 1 | NA07056 | 2 | - | 4th | - | 4th | 4th | 0.000 | 0.000 | 1.000 | 0.000 | 0.000 | 0.000 | 0.000 |
| 3 | NA06993 | 1 | NA07022 | 1 | - | 4th | - | 4th | 4th | 0.000 | 0.000 | 1.000 | 0.000 | 0.000 | 0.000 | 0.000 |
| 4 | NA07031 | 2 | NA12043 | 1 | - | 4th | - | - | 4th | 0.000 | 0.000 | 1.000 | 0.000 | 0.000 | 0.000 | 0.000 |
| 5 | NA12155 | 1 | NA12264 | 1 | - | 4th | - | 4th | 4th | 0.000 | 0.000 | 1.000 | 0.000 | 0.000 | 0.000 | 0.000 |
| 6 | NA12760 | 1 | NA12830 | 2 | - | FC | - | - | 4th | 0.000 | 0.000 | 1.000 | 0.000 | 0.000 | 0.000 | 0.000 |
| 7 | NA06989 | 2 | NA12155 | 1 | - | 4th | - | - | 5th | 0.000 | 0.999 | 0.001 | 0.000 | 0.000 | 0.000 | 0.000 |
| 8 | NA06991 | 2 | NA07022 | 1 | - | 4th | - | - | 5th | 0.000 | 0.997 | 0.003 | 0.000 | 0.000 | 0.000 | 0.000 |
| 9 | NA06994 | 1 | NA12892 | 2 | - | 4th | - | 5th | 5th | 0.000 | 1.000 | 0.000 | 0.000 | 0.000 | 0.000 | 0.000 |
| 10 | NA07014 | 2 | NA12043 | 1 | - | 4th | - | - | 5th | 0.000 | 1.000 | 0.000 | 0.000 | 0.000 | 0.000 | 0.000 |
| 11 | NA07031 | 2 | NA12761 | 2 | - | 4th | - | - | 5th | 0.000 | 0.994 | 0.006 | 0.000 | 0.000 | 0.000 | 0.000 |
| 12 | NA10831 | 2 | NA12264 | 1 | - | FCOR | - | - | 5th | 0.003 | 0.997 | 0.000 | 0.000 | 0.000 | 0.000 | 0.000 |
| 13 | NA10863 | 2 | NA12155 | 1 | - | FCOR | - | - | 5th | 0.000 | 1.000 | 0.000 | 0.000 | 0.000 | 0.000 | 0.000 |
| 14 | NA11931 | 2 | NA12748 | 1 | - | 4th | - | - | 5th | 0.000 | 1.000 | 0.000 | 0.000 | 0.000 | 0.000 | 0.000 |
| 15 | NA12752 | 1 | NA12830 | 2 | - | FCOR | - | - | 5th | 0.000 | 0.989 | 0.011 | 0.000 | 0.000 | 0.000 | 0.000 |
| 16 | NA12752 | 1 | NA12818 | 2 | - | 4th | - | - | 5th | 0.406 | 0.594 | 0.000 | 0.000 | 0.000 | 0.000 | 0.000 |
| 17 | NA12760 | 1 | NA12818 | 2 | - | FCOR | - | - | 5th | 0.000 | 1.000 | 0.000 | 0.000 | 0.000 | 0.000 | 0.000 |

**Figure 7:**
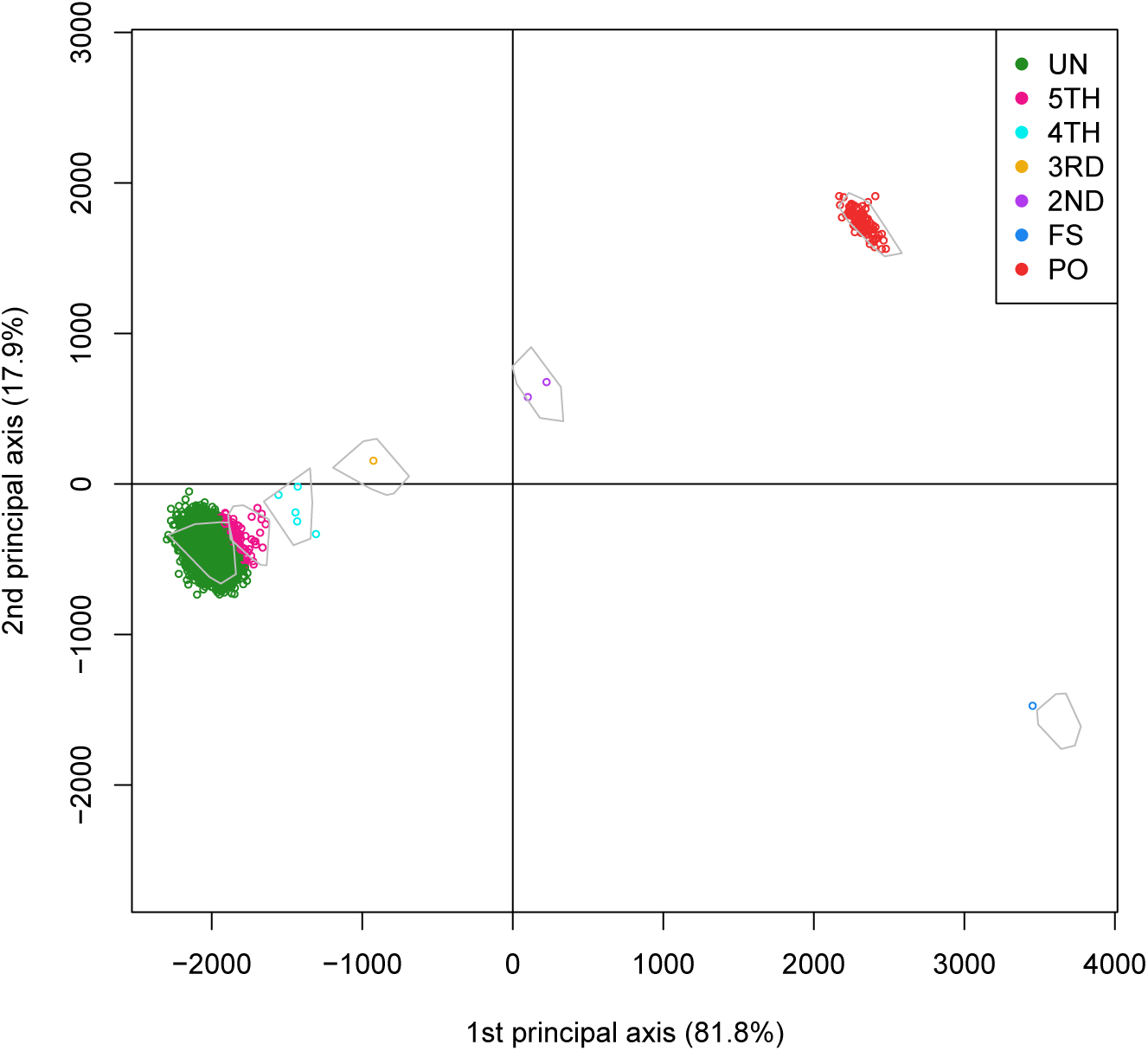
MDS map of 13,530 pairs of individuals of the CEU population. Convex hulls delimit the region of the pairs obtained by genetic bootstrapping.

Our results confirm a third degree pair (pair 1 in Table 4) reported by Kyriazopoulou *et al.* (2011). We also confirm four of the fourth degree pairs reported by the latter authors (pairs 2-5 in Table 4). However, we also observed considerably incongruence of our results with those of the latter authors. We found an FC pair to be classified as fourth degree (pair 6) by our method and 11 reported fourth degree pairs were classified as fifth degree. One of these (pair 16) had a high probability of being even more remotely related. We also compared results with those published by Huff *et al.* (2011), who estimate recent shared ancestry (ERSA) by using IBD segments. Our work confirms three fourth degree pairs and one fifth degree pair report by the latter authors, though we found two additional fourth degree pairs which are not confirmed by Huff *et al.* (2011).

### 4.2 GCAT samples

We use samples from the GCAT Genomes for life project, a cohort study of the genomes of Catalonia (www.genomesforlife.com). GCAT is a prospective cohort study that includes 17,924 participants (40-65 years, release August 2017) recruited from the general population of Catalonia, a Mediterranean region in the northeast of Spain. Participants are mainly part of the Blood and Tissue Bank (BST), a public agency of the Catalan Department of Health. Detailed information regarding the GCAT project is described in Obo´n-Santacana *et al.* (2018). We study relatedness on 5,075 GCAT Spanish participants from Caucasian origin and 736,223 SNPs that passed quality control [Sumoy et al., 2017]. Inferred relatives of first and second degree were confirmed by the BST public agency, for pairs sharing one surname (PO, second degree pairs) or two surnames (FS pairs), respecting the privacy of the participants. According to the same filtering procedures used in the CEU samples, 26,006 SNPs (MAF *>* 0.40, LD-pruned, HWE exact mid p-value *>* 0.05, and missing call rate 0) were considered for relatedness analysis. A MDS map of the individuals, using the allele sharing distance, is shown in Figure 8A.

**Figure 8:**
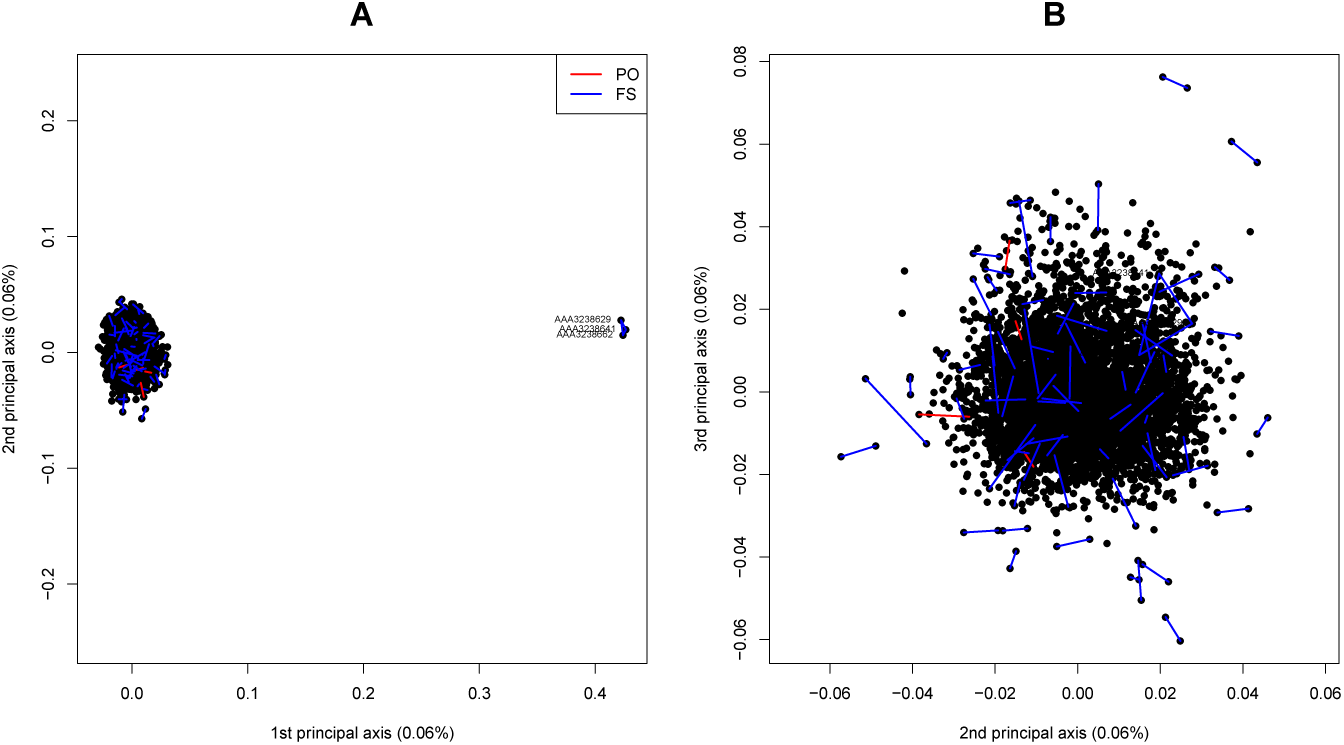
MDS maps of 5,075 individuals of the GCAT study.

The first dimension shows an outlying triple of sibs (pairs 48, 49 and 50 in Table S1, Figure 8A). When considering second and third dimensions, this trio is masked, and these axes reveal other FS pairs (Figure 8B). Figure 9 shows the IBS-MDS approach for pairs of individuals using the Euclidean distance. Our IBS-MDS method detects four PO, 70 FS and 12 second degree pairs, and reveals 68 and 54 third and fourth degree pairs (See supplementary Table S1). The IBS-MDS-plot shows a large variability in the FS group, where many pairs fall outside their respective simulated convex hull. We suggest this excess variability is, at least in part, due to the existence of some three-quarter siblings in the database (e.g. half-sibs where two of the three parents have an FS or PO relationship). The Cotterman coefficients for a three-quarter sibling relationship are readily deduced to be *k*_0_ = 3*/*8, *k*_1_ = 1*/*2 and *k*_2_ = 1*/*8, and their co-ancestry coefficient is *θ* = 3*/*8, or, equivalently, their kinship coefficient is 3*/*16. They can therefore be expected to fall in-between the

**Figure 9:**
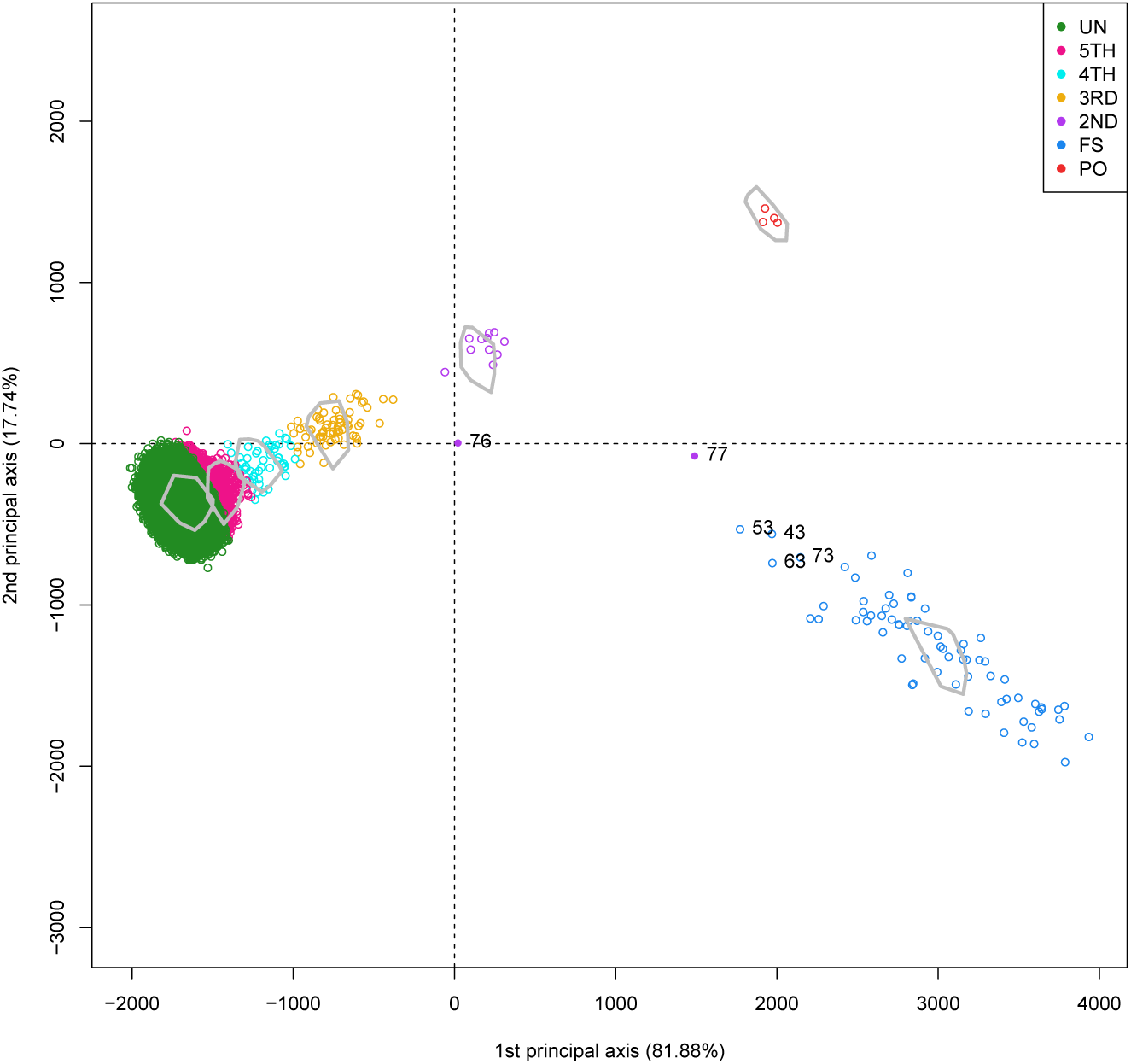
MDS map of pairs of individuals of the GCAT study. Convex hulls delimit the region of the pairs obtained by genetic bootstrapping.

FS and 2nd degree clusters, as is shown for some labeled pairs in Figure 9. We therefore repeated IBS-MDS with a new training set where this type of relationship had been included. The resulting map is shown in Figure 10 and shows that one outlying second degree pair (77) and several initially outlying FS pairs (73, 63, 53, 43) are now indeed classified as three-quarter siblings. The posterior probabilities for these classifications are given in supplementary Table S2.

**Figure 10:**
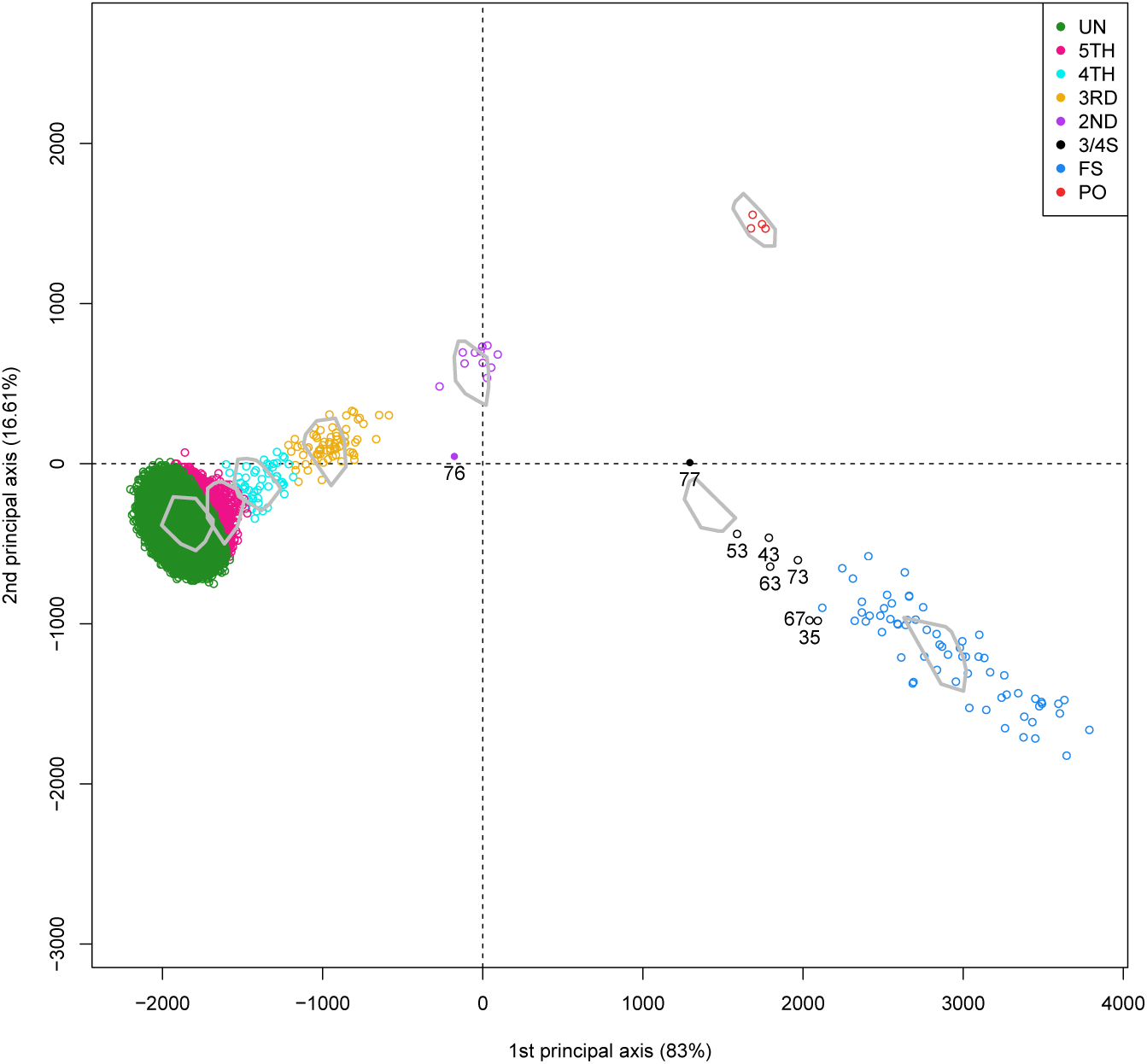
MDS map of pairs of individuals of the GCAT study, trained with an additional category for three-quarter siblings (3/4S). Convex hulls delimit the region of the pairs obtained by genetic bootstrapping.

## 5 Conclusions and Discussion

Multidimensional scaling is a widely used tool in genetic analysis, and is often used to check data for homogeneity and for substructure investigation. If there exists population substructure, MDS maps typically reveal clusters corresponding to the different populations. However, outlying observations or groups of outlying observations do not necessarily *prove* the existence of population substructure. Individuals may separate out in MDS because they are related but actually stem from the *same* population. We conclude that population substructure cannot be ruled out or confirmed only by looking at MDS maps. We consider it necessary to look for additional zero eigenvalues and to inspect diagnostic plots like Figure S1 to identify influential observations, and to perform a relatedness study before inferring the existence of any population substructure.

The proposed method for classifying pairs using MDS and discriminant analysis is seen to perform well with both simulated and empirical data. The sampling of artificially related pairs from the observed data requires a considerable number of approximately unrelated individuals to be present. We therefore suggest the method to be used for large samples, where such a substantial subset of unrelated individuals can be identified. This is probably not an obstacle for the use of our method, as increasingly large samples are being used in epidemiological genomics. The sampling of artificial pairs from the observed data respects the allele frequency distribution of the original data, and provide reference areas for the different relationships given the allele frequencies of the observed data.

If MDS is used with the primary purpose of detecting substructure, we suggest first to remove one individual from each related pair, and to apply MDS to the filtered dataset. If not, related individuals can upset the proper detection of substructure. Incidental outliers in MDS maps are likely to correspond to individuals that have a close family relationship. Figure 2 shows a set of standard patterns that are relatively easy to recognize in genetic data analysis. However, if more family relationships are present in the database, the patterns obtained by MDS are more difficult to interpret.

The proposed IBS-MDS method can be used in an exploratory and iterative way. For some data sets, a first analysis may reveal that certain relationship categories are obviously absent in the data. In such cases, IBS-MDS can be re-run with a training set without these categories, in order to focus the analysis on those relationships that are of relevance for the data under study. Indeed, we tacitly omitted monozygotic twins (MZ) from our training set, as MZ pairs (or duplicates) are easily recognized, and this way the analysis is focused on more remote relationships. The GCAT example illustrates this iterative use, as three-quarter siblings were added to the training set in second instance to improve the analysis.

In standard applications of MDS to bi-allelic genetic variants, genetic data is often coded in (0,1,2) format and Euclidean distances between individuals are calculated in this coding, and represented by MDS in low-dimensional space. The units on the axes of such maps are not very well interpretable, because the Euclidean distances are calculated over a large number of genetic variants. We propose to scale the Euclidean distance matrix that is entered into MDS by a factor of 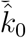, where *k* is the number of variants. This scaling will not affect the configuration of the points in the map, and neither its goodness-of-fit, but it will render the axes and distances of the map more interpretable. Two individuals that are a unit distance apart in the map now differ on average by one allele per locus. This will typically scale all the coordinates in the MDS map within the (−1,1) interval, as the maximum difference in number of alleles between two individuals is two. We note that the Euclidean distance is not the same as the allele-sharing distance we used in this paper. The allele-sharing distance is the sum of the absolute value of the difference in numbers of alleles over all loci. In multivariate analysis this metric is better known as the Manhattan distance, the city-block distance, or the taxicab metric. Division of the Manhattan distance by the number of polymorphisms gives the allele-sharing distance. For the sake of interpretability, it is convenient to scale the Manhattan distance matrix by 1*/k* prior to using it in MDS. A pair of individuals two units apart will then differ by two alleles at all loci. The suggested scalings facilitate interpretation of MDS results, but are hampered by the fact that the map is a low-dimensional approximation to the original distance matrix, and therefore the property will not hold exactly, but only approximately so. For distance matrices that have the Euclidean property, and that therefore only have non-negative eigenvalues [Mardia et al., 1979, Chapter 14], the map distances will approximate the true distances from below, and one can say that two individuals that are, for instance, 0.5 units apart, will differ on average by at least 0.5 minor alleles per locus, or can be expected to have at least one different allele for every pair of loci. We have used Manhattan distances scaled by 1*/k* in all MDS maps of individuals used this paper (Figures 1, 3, 6 and 8).

We note that the family relationships detected in Figure 3 would have gone largely unnoticed if the data is analyzed by non-metric multidimensional scaling of the between-individual distance matrix. In non-metric MDS only the rank order of the distances is used, not the value of the distances itself. In non-metric MDS related individuals do therefore generally not appear as outliers. This makes the non-metric approach less attractive at the individual level, though it does not rule out the non-metric approach for analysing pairs.

We recommend the use of discriminant analysis in allele-sharing studies as employed in this paper. The posterior probabilities of the different relationships give a quantitative criterion for deciding upon which relationship is most likely for a given pair of individuals. In allele sharing studies this decision is mostly made graphically by inspecting a (*p*_0_*, p*_2_) plot in IBS studies, or a (*k*_0_*, k*_2_) plot in IBD studies. We note that these posterior probabilities differ from those used in a standard discriminant analysis, in the sense that they are affected by additional uncertainty generated by using a training set obtained by genetic bootstrapping of the observed data.

The GCAT sample shows, for almost all relationship categories, larger variability in the MDS coordinates than would be expected under strict Mendelian sampling of alleles from unrelated individuals. This excess variability can, at least in part, be explained by the presence of additional relatedness between (unobserved) close relatives of the individuals in the database. This leads to increased autozygosity, which is a characteristic of more endogamous populations. The occurrence of three-quarter siblings is just a particular instance of this phenomenon. Consequently, the degree of relatedness of two individuals tends to become a continuous variable, which is increasingly hard to discretize into the standard relationship categories.

Several scholars have recently distinguished between remote or background relatedness (population substructure) and more recent relatedness (family relationships), describing methods that try to address both [Conomos et al., 2016, Weir and Goudet, 2017]. We have shown in this paper that MDS can be used to detect both recent relatedness and substructure.

## Acknowledgments

This work was partially supported by grants SGR551 and SGR1269 from the Agència de Gestio´ d’Ajuts Universitaris i de Recerca (AGAUR) of the Generalitat de Catalunya, by grant MTM2015-65016-C2-2-R and ADE 10/00026 (MINECO/FEDER) of the Spanish Ministry of Economy and Competitiveness and European Regional Development Fund, and by grant R01 GM075091 from the United States National Institutes of Health. R. de Cid was supported by the Ramon y Cajal action (RYC-2011-07822).

---

**Figure S1:**
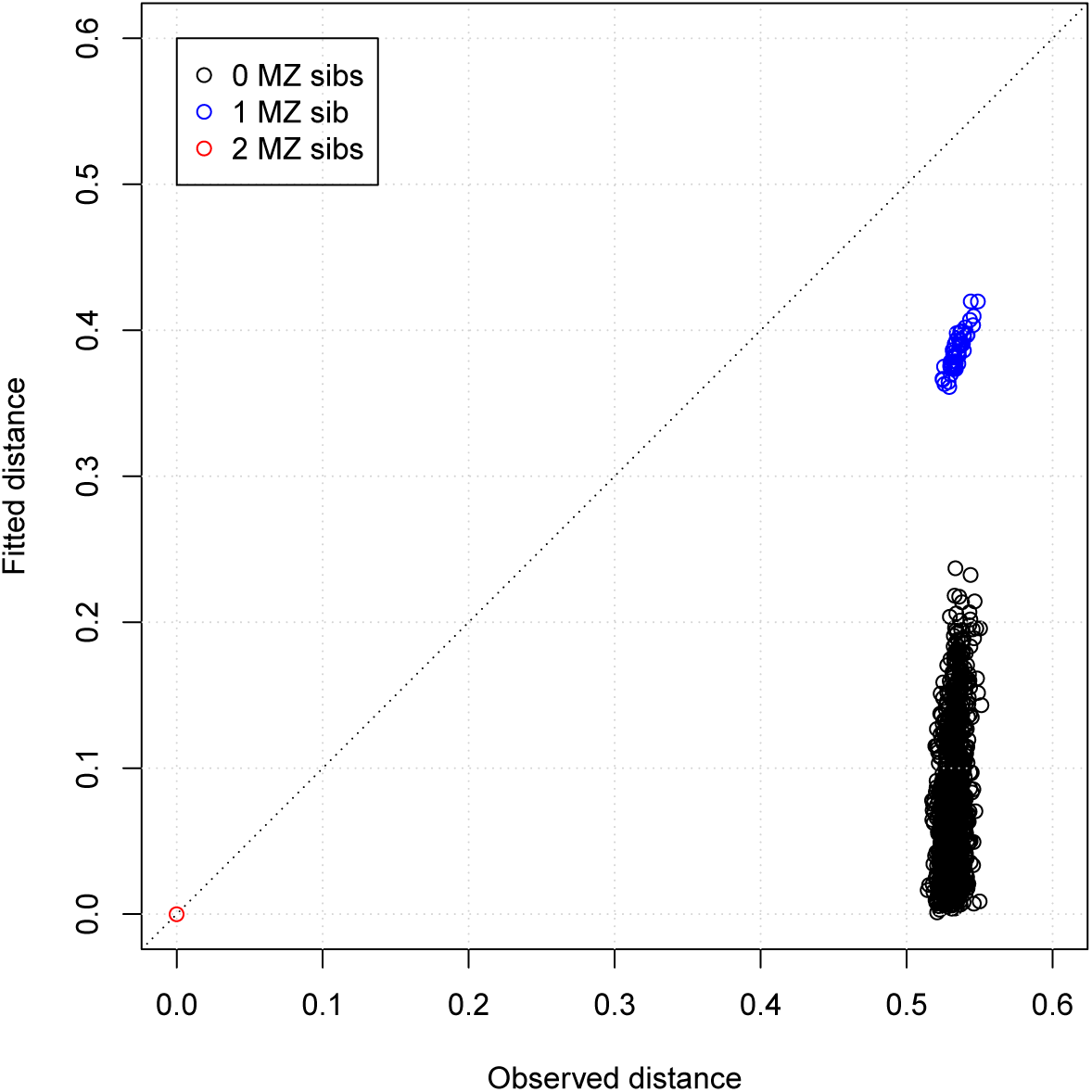
Fitted versus observed distance for an MDS map of a homogeneous population with one MZ pair. The single pair involving the two individuals of the MZ pair is red, pairs involving one individual of the MZ pair are blue and pairs not involving any individual of the MZ pair are black.

**Figure S2:**
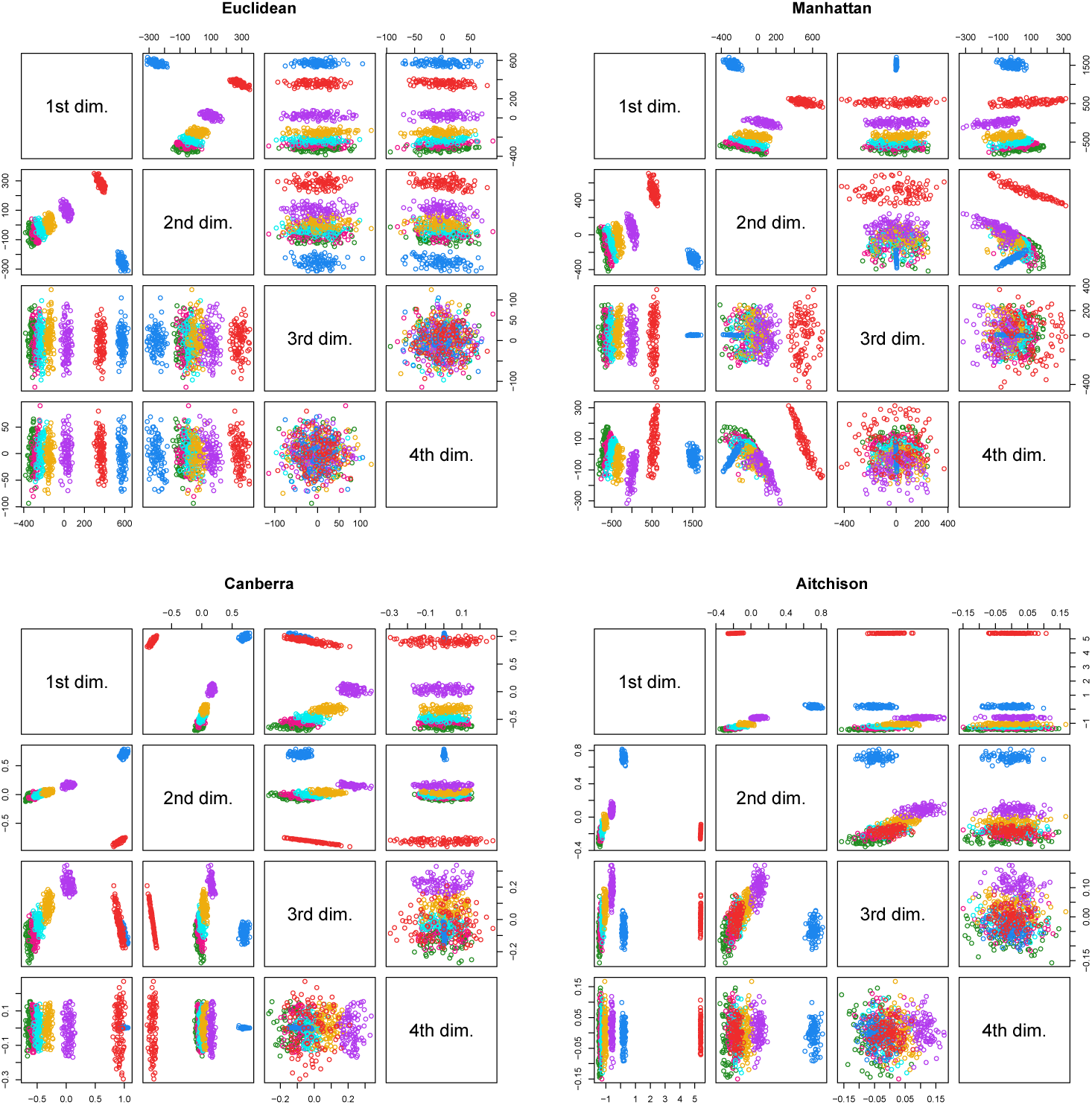
Scatterplot matrices of the coordinates of the first four dimensions of an MDS solution using Euclidean, Manhattan, Canberra and Aitchison distance. Data obtained by simulation of 5,000 independent variants with a MAF of 0.5 under Hardy-Weinberg equilibrium.

**Table S1:**
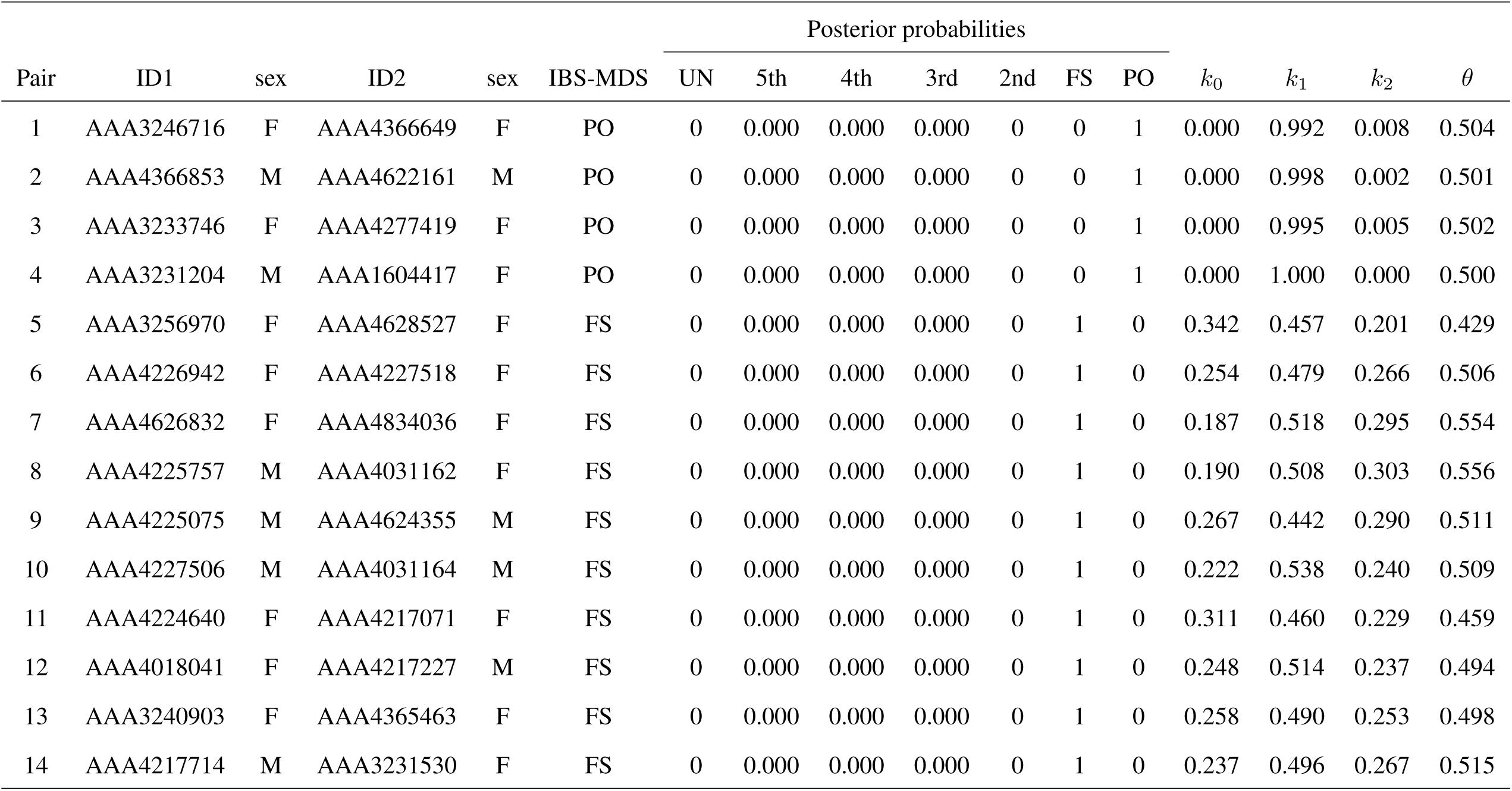
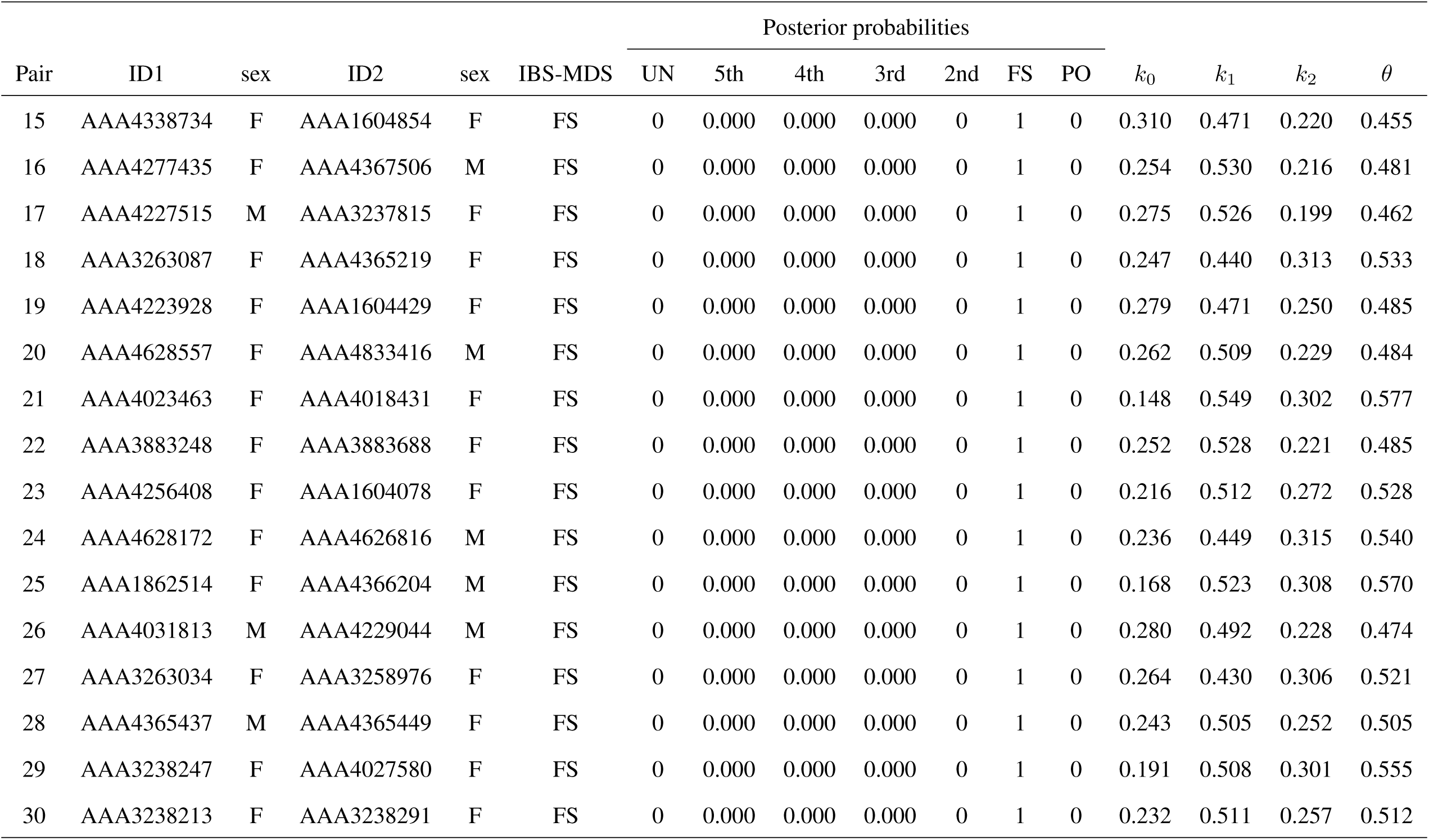
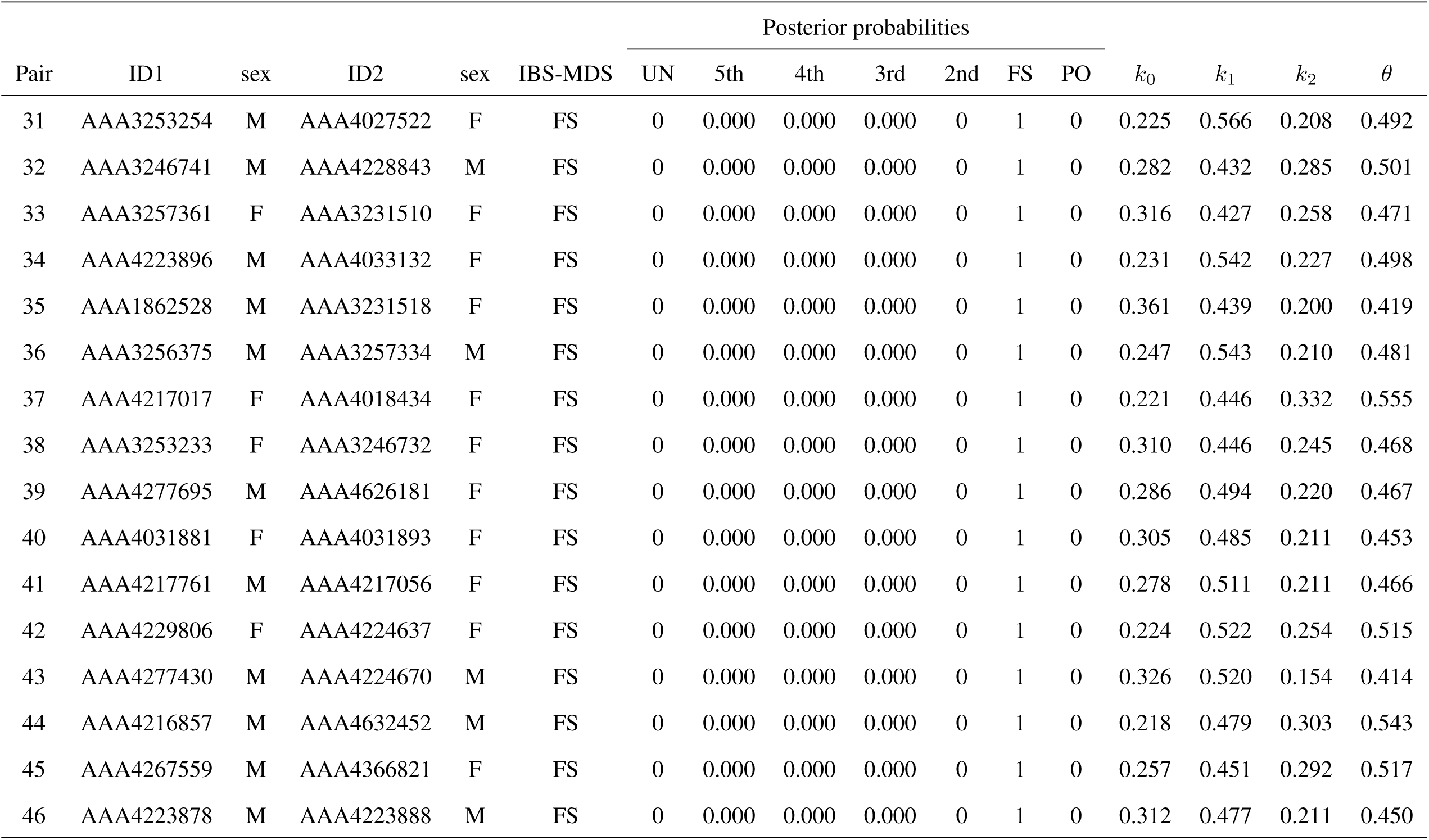
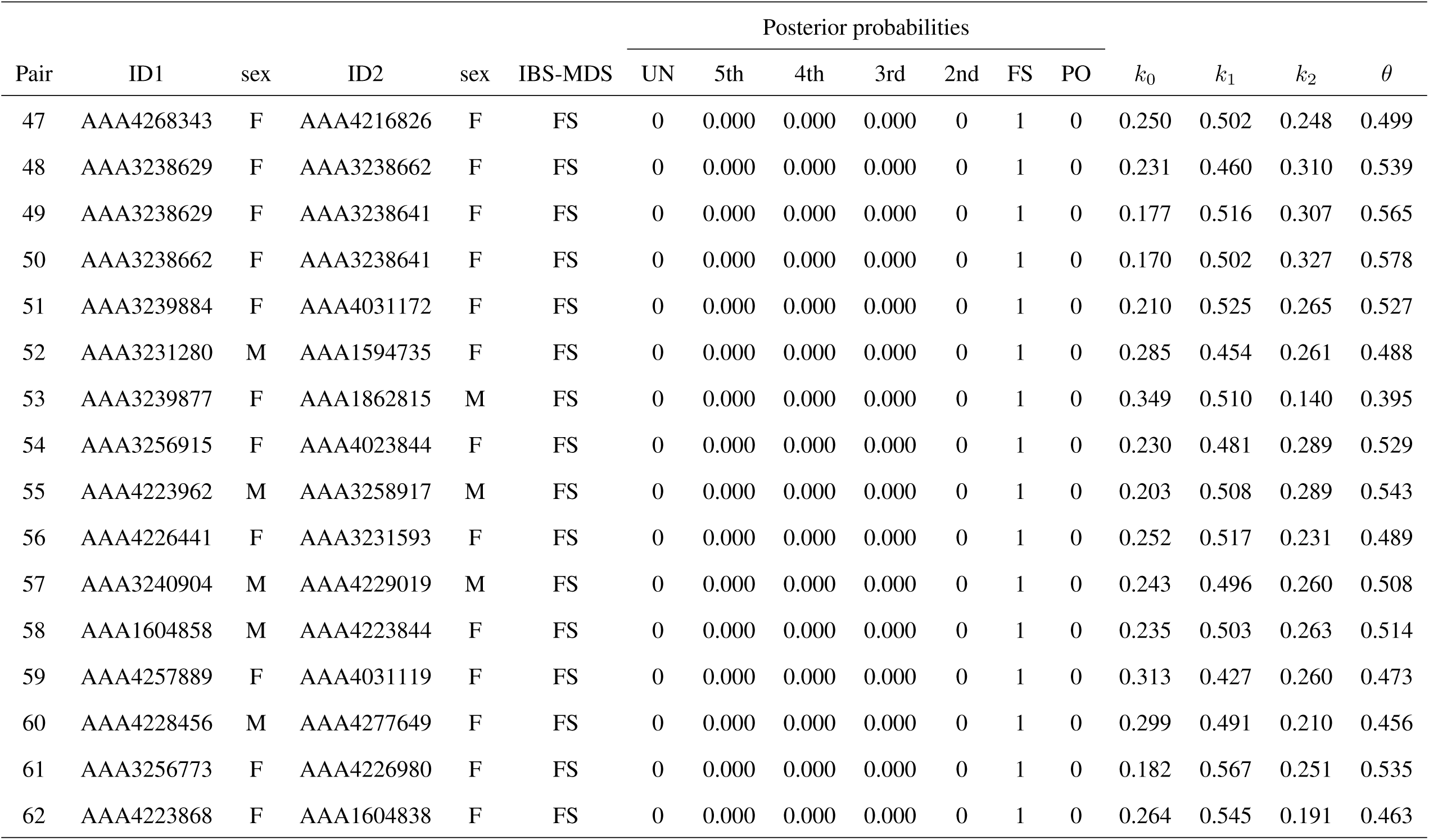
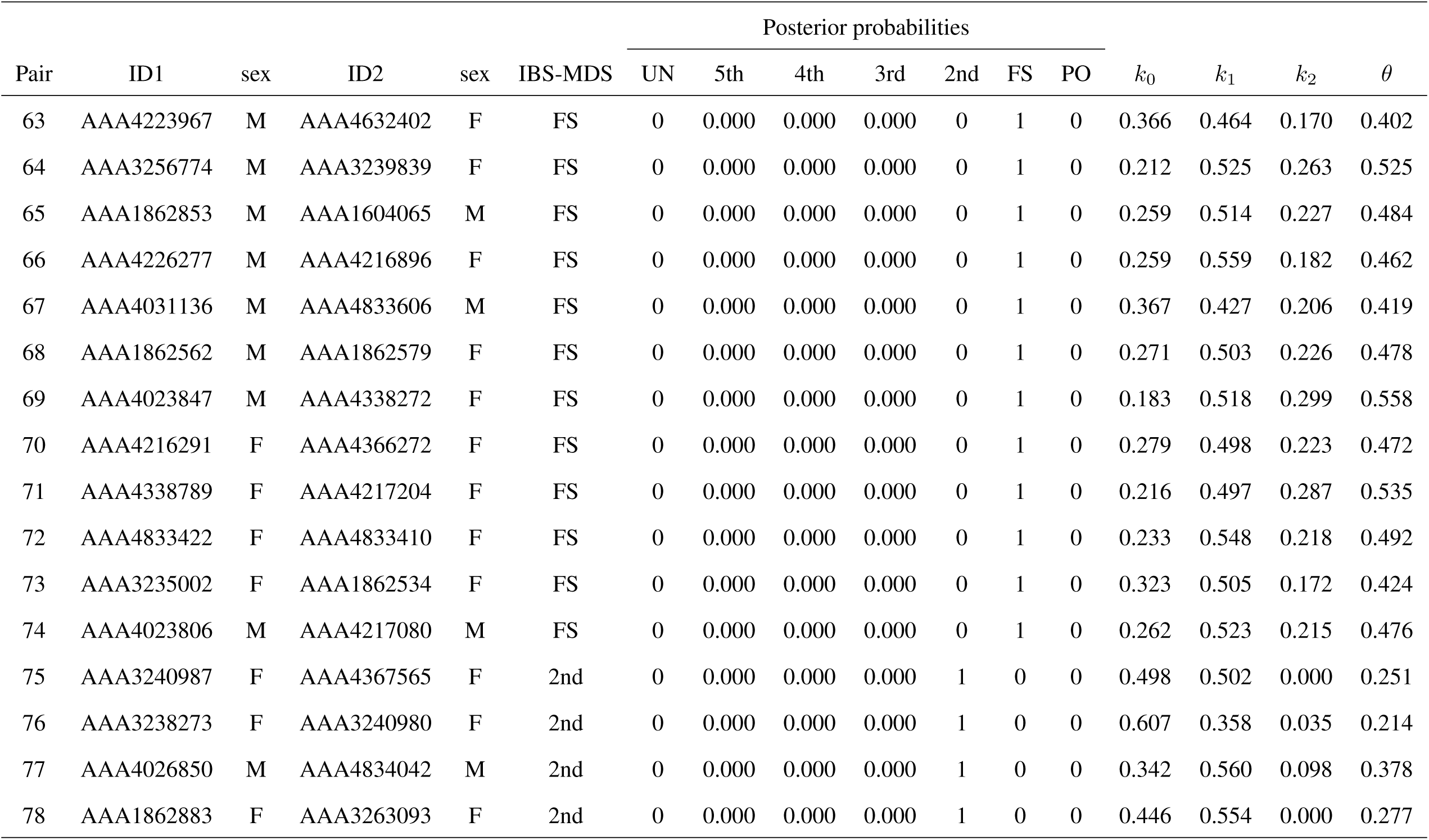
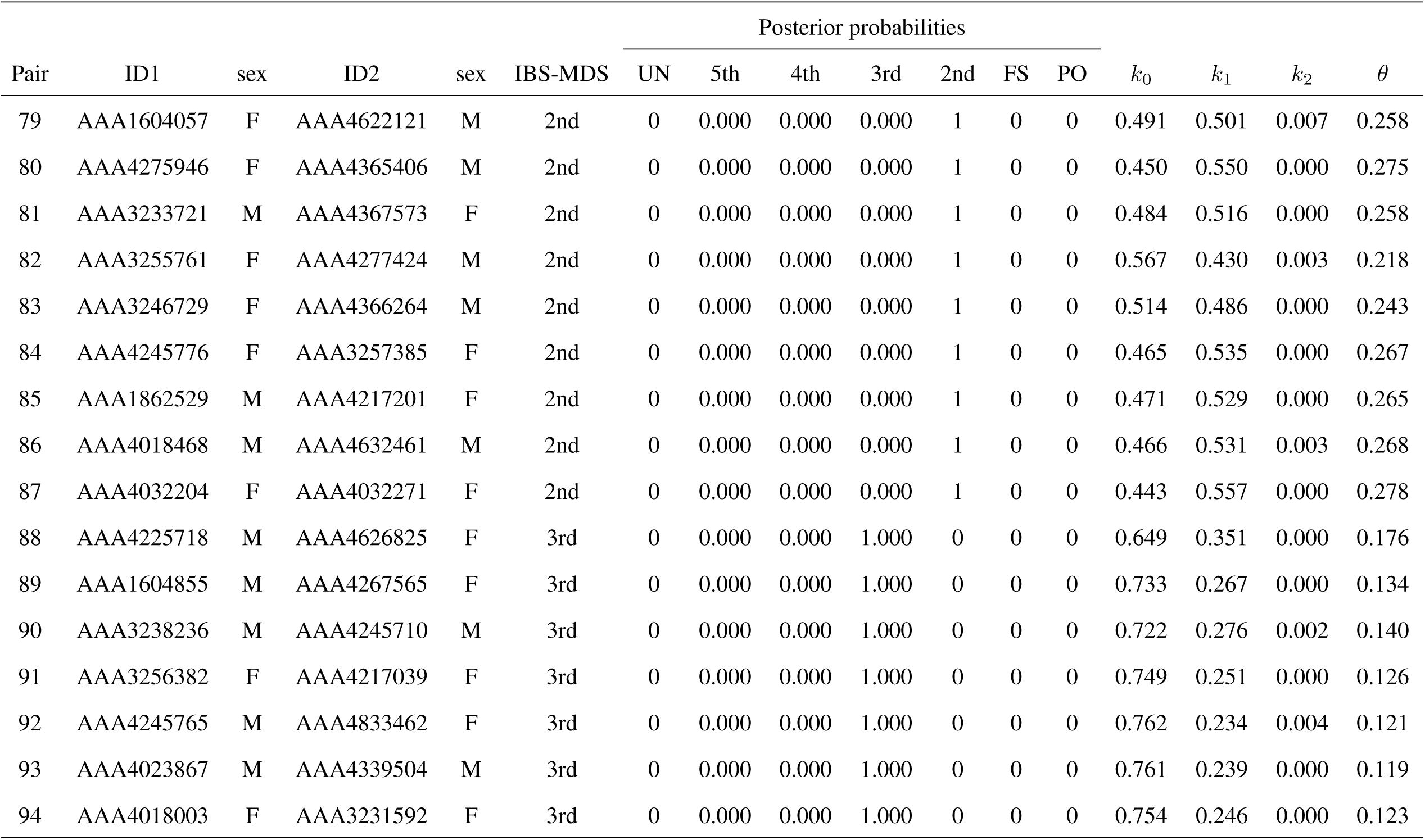
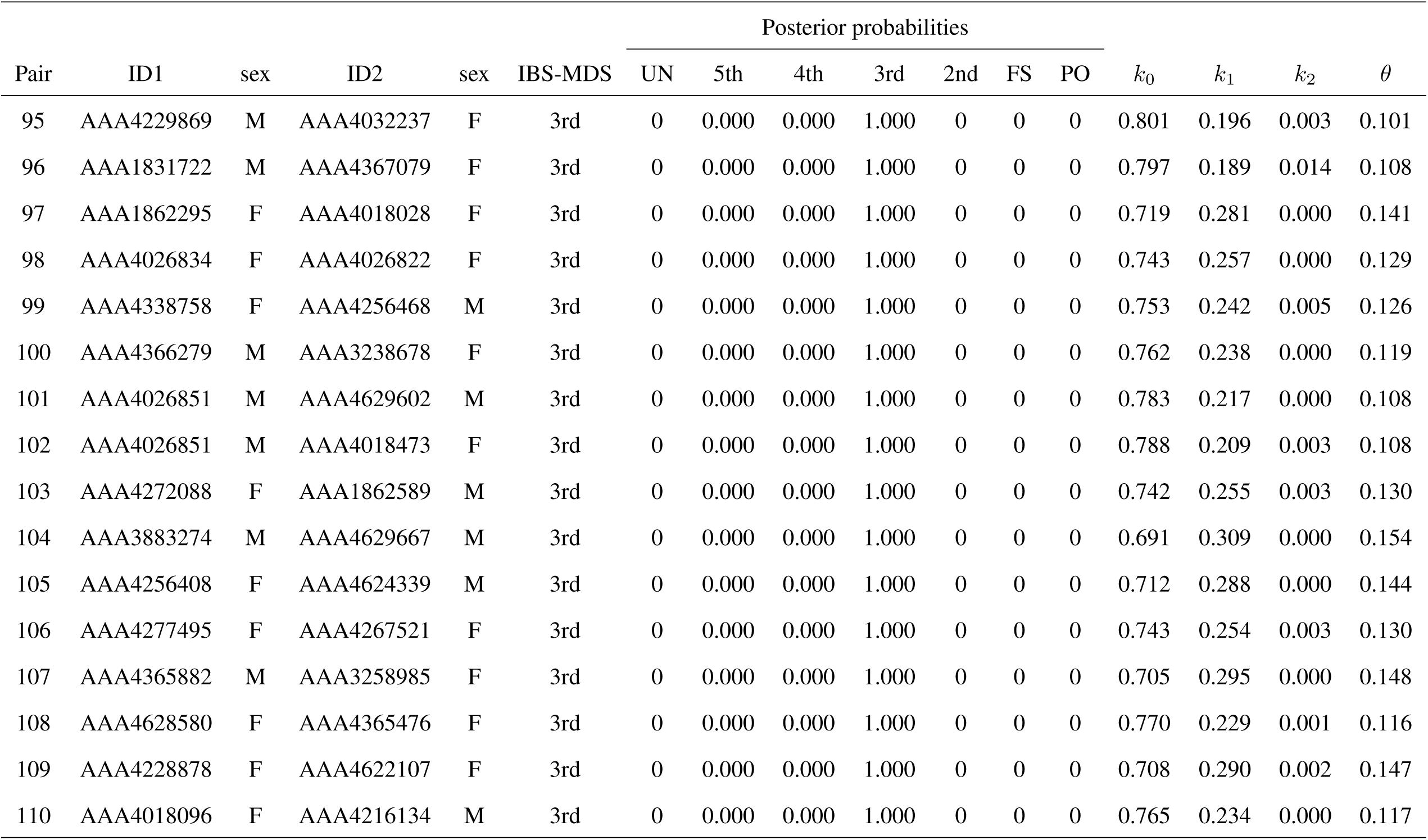
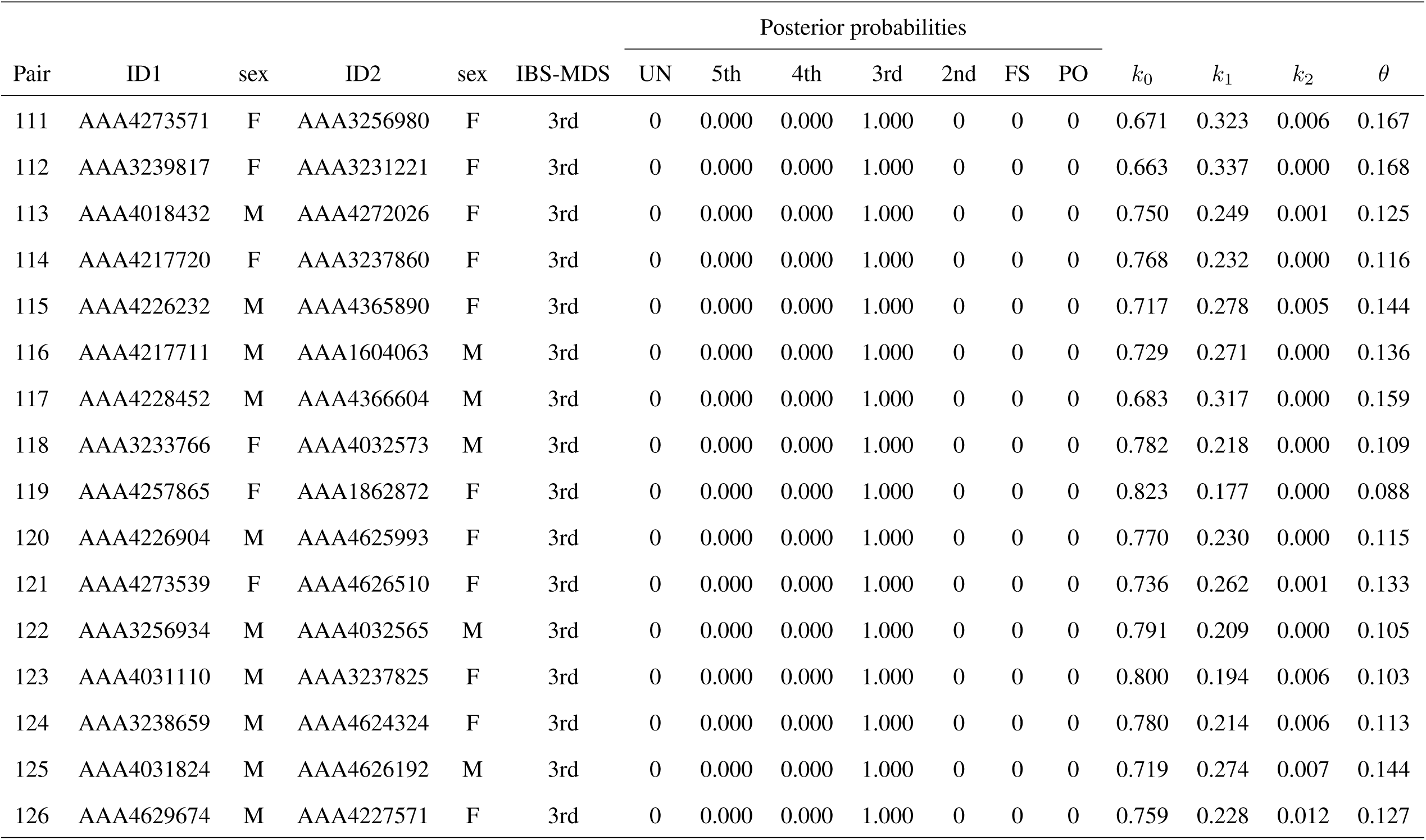
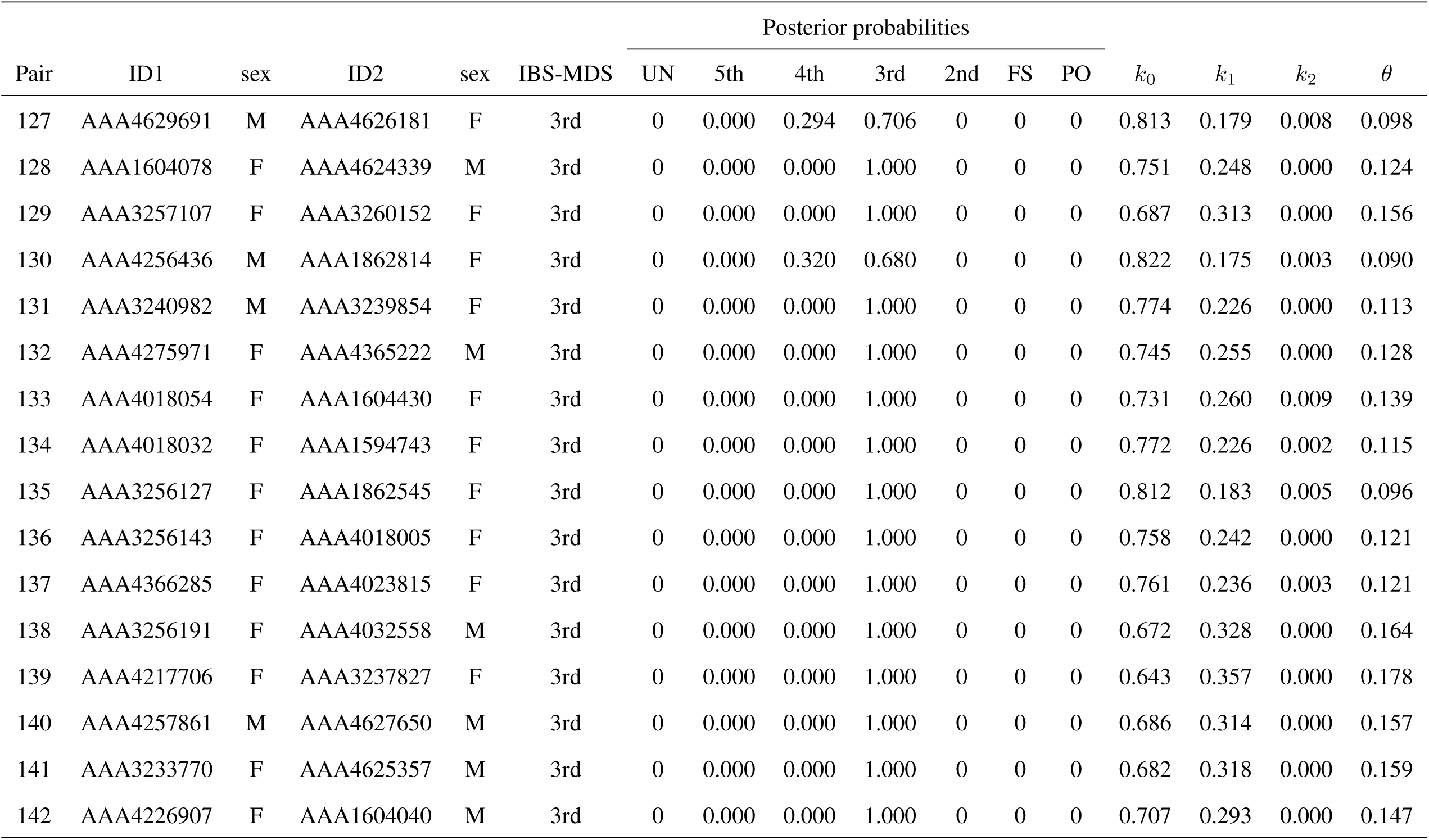
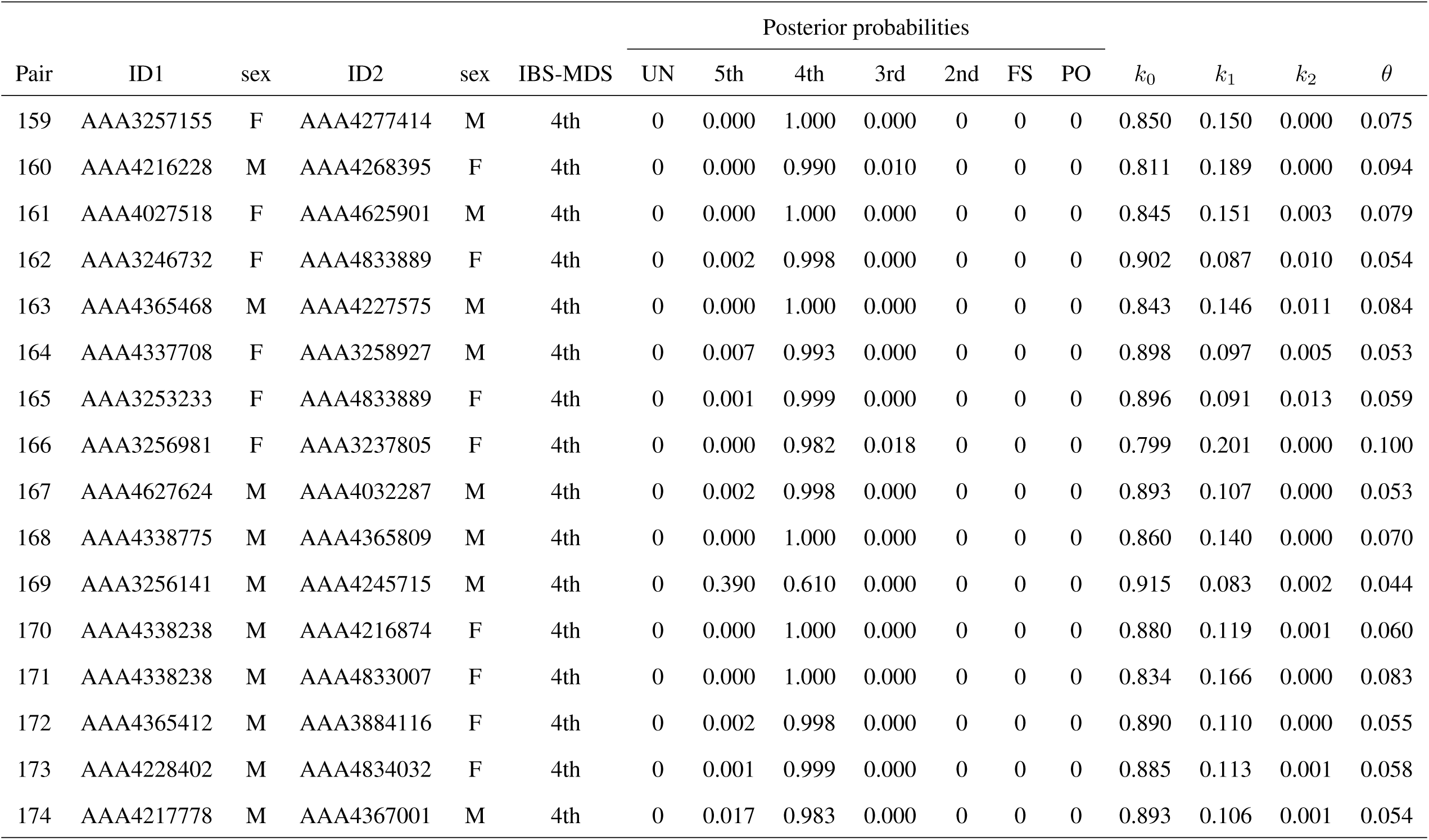
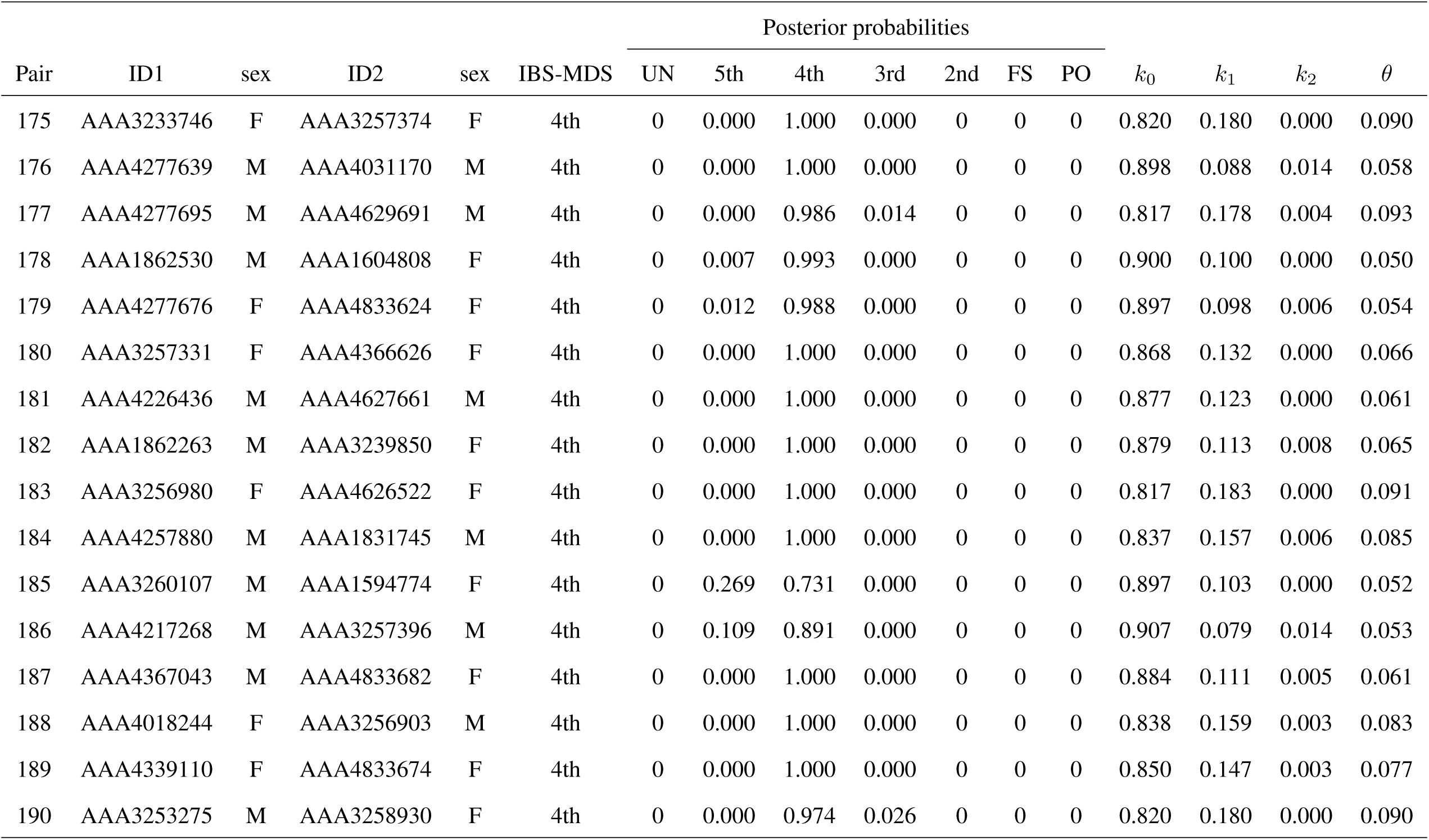
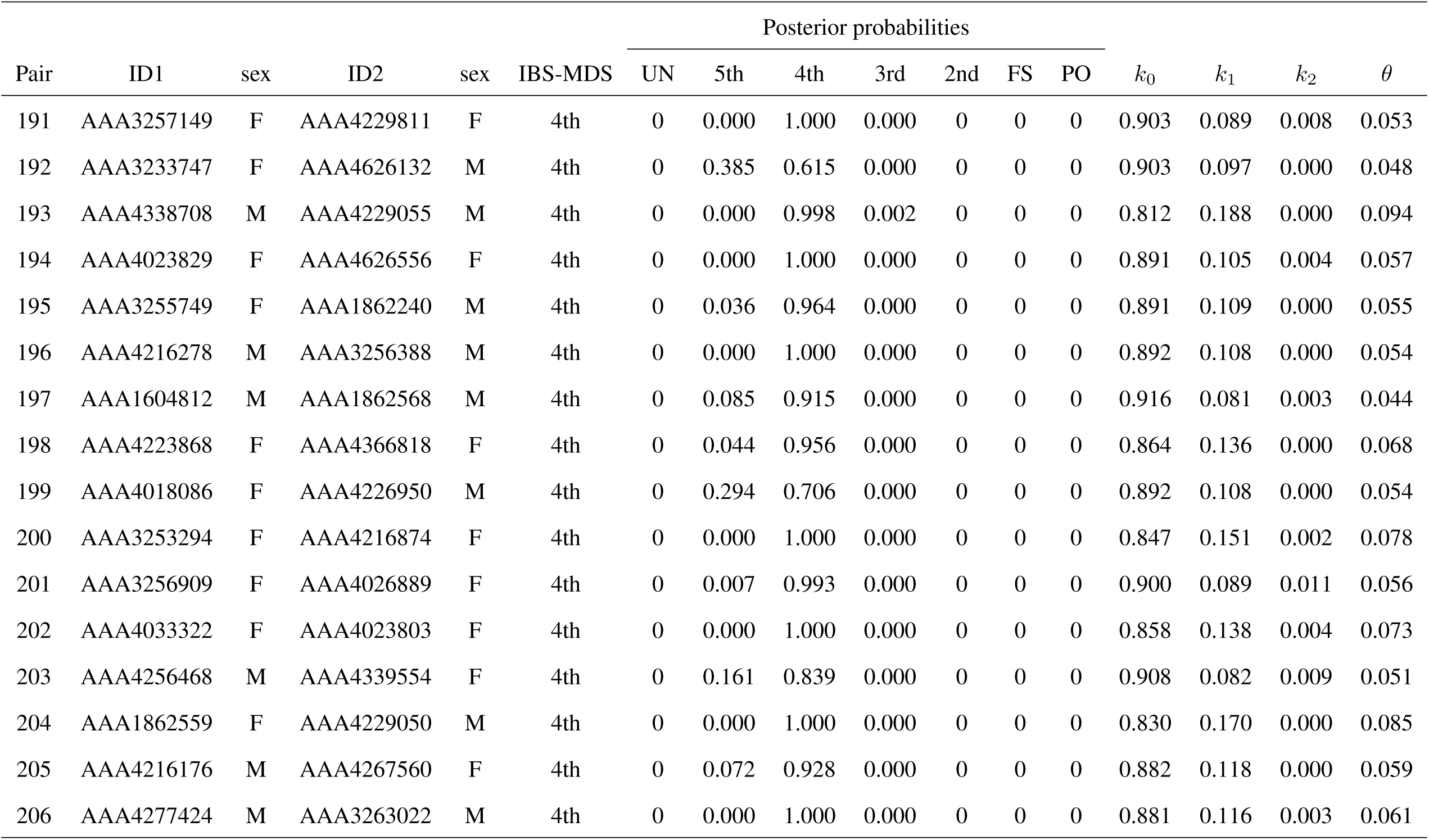
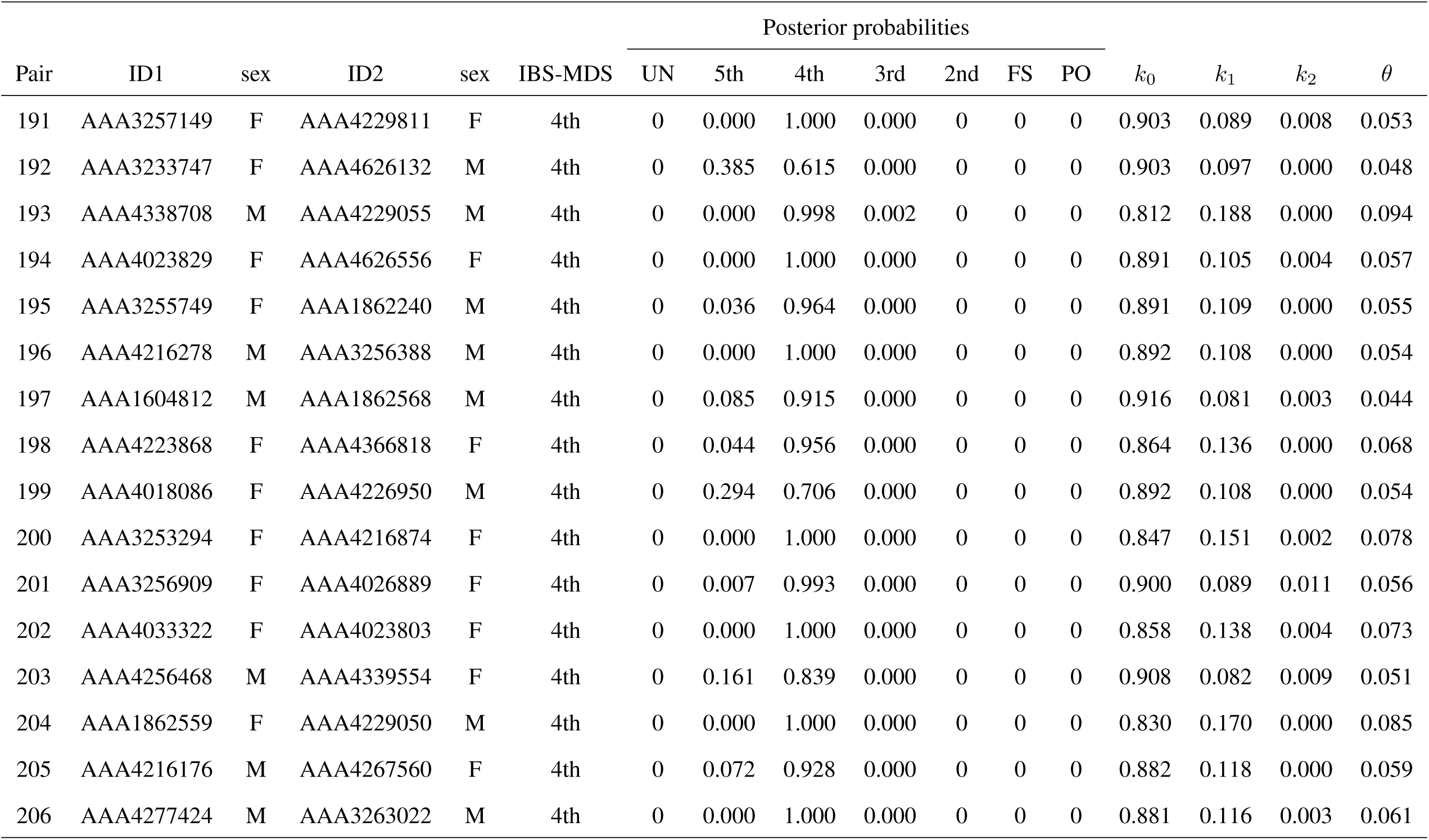
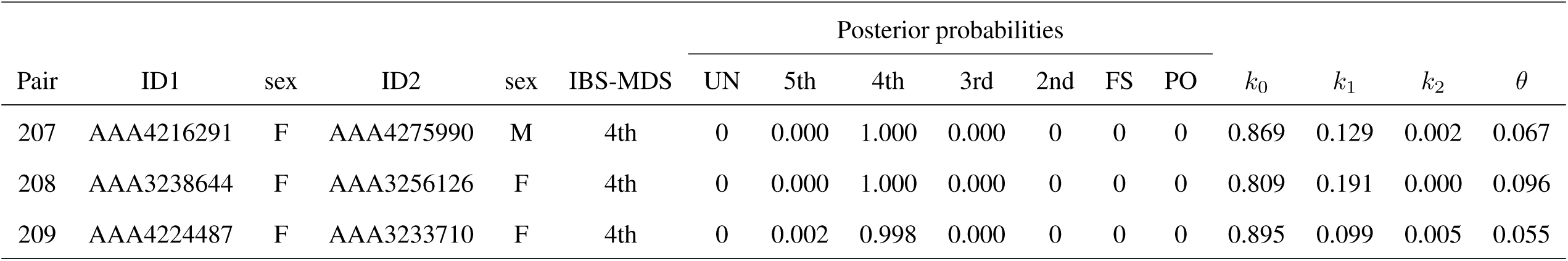
Predicted relationships of first, second, third and fourth degree pairs of the GCAT cohort and their probabilities according to IBS-MDS. *k*0, *k*1, *k*2: Cotterman’s coefficients, *θ*: coancestry coefficient.

**Table S2:**
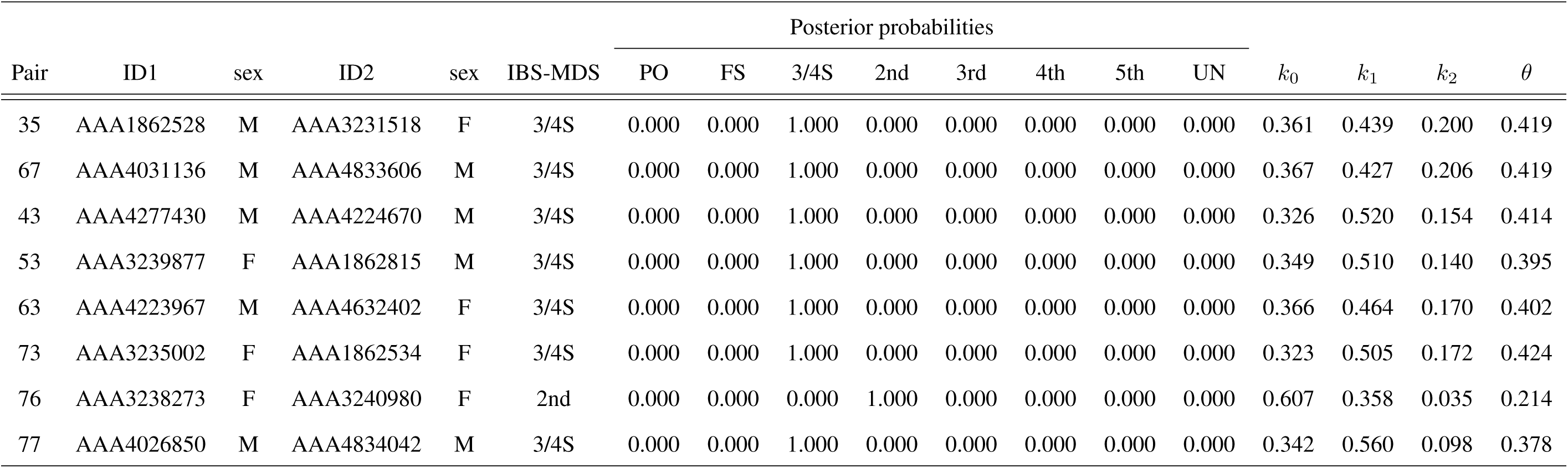
Potential three-quarter siblings of the GCAT cohort and their probabilities according to IBS-MDS. *k*0, *k*1, *k*2: Cotterman’s coefficients, *θ*: coancestry coefficient.

